# PARP inhibition enhances exemestane efficacy in triple-negative breast cancer

**DOI:** 10.1101/2024.07.31.605956

**Authors:** Nur Aininie Yusoh, Liping Su, Suet Lin Chia, Xiaohe Tian, Haslina Ahmad, Martin R. Gill

## Abstract

Triple negative breast cancer (TNBC) remains the breast cancer subtype with the poorest prognosis and median survival rate. Targeting PARP1/2 with PARP inhibitors (PARPi) and achieving synthetic lethality is an effective strategy for TNBCs with BRCA1/2 mutations, however, the majority of TNBCs are BRCA1/2 wild type. Synergistic drug combinations with PARPi offers the potential to expand the use of PARPi towards BRCA-proficient cancers, including TNBC. To identify new PARPi combinations, we screened a library of 166 FDA-approved oncology drugs for synergy with the PARPi Olaparib in TNBC cells. We found that Exemestane, an aromatase inhibitor, synergised with Olaparib with a significant decrease in IC_50_ values and clonogenicity accompanied by elevated DNA damage and apoptosis seen in combination treatment. The mechanistic basis for synergy was rationalised by the previously unreported ability of Exemestane to induce replication stress, as evidenced by ATR pathway activation and RPA foci formation. Low impact of this combination towards normal breast epithelial cells was observed and Exemestane has no reported severe toxicity as a monotherapy. This combination was able to achieve enhanced tumor growth inhibition in a murine xenograft model, greater than either drug employed as a single-agent. GO and KEGG enrichment analysis of differential genes indicated alterations in pathways associated with cell death in response to Exemestane and Olaparib treatment.

## INTRODUCTION

Comprising 15-20% of all breast cancer cases [1], triple-negative breast cancer (TNBC) is an aggressive subtype characterized by the lack of expression of estrogen receptor (ER), progesterone receptor (PR), and human epidermal growth factor receptor type 2 (HER2). Whilst non-TNBC patients have benefited from major improvements in personalized therapy such as estrogen therapy or HER2-targeted treatment, only a small number of novel therapeutic agents have gained approval for TNBC [2]. These limited treatment options combined with the heterogeneous nature and high metastatic potential of TNBC often leads to early relapse and a poorer prognosis compared to other subtypes [3].

A recent advancement in targeted therapeutics are poly(ADP-ribose) polymerase (PARP) inhibitors (PARPi), which have shown promise as anti-cancer therapeutics, particularly towards BRCA-mutated BC [4–6]. Mechanistically, this relies on the principle that when PARP enzymatic activities are inhibited, DNA single-strand breaks (SSBs) are converted to DNA double-strand breaks (DSBs). If the homologous recombination (HR) repair pathway is deficient, such as in those patients with BRCA mutations, DSBs remain unrepaired, leading to genome instability and/or cell death. However, in BRCA-proficient cells, these DSBs damage are repaired via HR pathway and thus, PARP inhibition alone may not necessarily induce cell death. Hence, the efficacy of PARPi monotherapy is limited to BRCA-mutated BC patients and the majority of TNBC, particularly those without BRCA1/2 mutations (80-90% of TNBC patients) [7, 8], still do not benefit from PARPi.

Drug combinations employing synergistic pairs can achieve improved anti-tumor responses with lower concentrations of single agents and overcome resistance [9, 10]. For PARPis, rational combination selection focussing on complementary mechanisms have attracted considerable interest as a potential method by which to expand the scope of PARPis, including towards BRCA-proficient cancers [11]. This has seen numerous DNA-damaging agents achieve synergy with PARPi, including platinum drugs [12, 13], doxorubicin [14], and gemcitabine [15], whilst targeting alternative DNA repair pathways has also achieved success in this capacity [16]. An alternative method to identify drug combinations is unbiased (or hypothesis-free) screening [17]. By utilizing chemical libraries alongside the drug of interest, this has the advantage of allowing for serendipitous discoveries and can facilitate greater understanding of the polypharmacology of drugs, where subsequent target deconvolution studies of a newly identified combination can reveal novel, sometimes unexpected, mechanisms of action [18].

Moreover, when an FDA-approved library is employed, clinical translation may be accelerated. Despite this, such unbiased synergy screens involving PARPi are relatively rare. The validity of this approach may be demonstrated by Lui et al., who reported the high-throughput drug combination screening of PARPi rucaparib alongside a compound library consisting of 395 FDA-approved and investigational compounds [19]. Strikingly, the authors found that the effect of Bromodomain and Extra-Terminal motif (BET) inhibitors was enhanced by rucaparib in patient-derived ovarian cancer cells and, encouragingly, that this effect occurred *irrespective* of homologous recombination deficiencies.

In this study, we performed an unbiased drug synergy screen employing Olaparib, the most successful PARPi to date, alongside an FDA-approved oncology drug library to identify synergistic combinations. We uncovered that Exemestane and Olaparib are synergistic in BRCA-proficient TNBC cells, likely due to the hitherto unknown ability of Exemestane to generate replication stress, characterize the mechanism of action of drug synergy and validate this combination in a murine model. Altogether, our study provides a novel combination of Olaparib active towards TNBC *in vitro* and *in vivo* and reveals a new mechanistic function of Exemestane.

## RESULTS

### Exemestane shows strong synergism with Olaparib in TNBC cells

First, a drug combination screen was performed employing an FDA-approved oncology drug library of 166 drugs (details within Supplementary Table S1) and a non-cytotoxic concentration of Olaparib in MDA-MB-231 TNBC cells (10 μM Olaparib; 24h IC_50_ of Olaparib > 100 μM, Supplementary Fig. S1), as depicted in Fig. 1. Resultant cell viabilities following treatment were determined by MTT assay and compared to single-agent treatment conditions. Chou and Talalay combination index (CI) analysis were performed to determine synergy [20, 21]. This identified ten compounds (16.9% of all drugs tested) that gave more than two synergistic CIs (CI < 0.9) when combined with Olaparib over the range of concentrations tested (Supplementary Table S2), with substantial enhancement in cytotoxicity profiles than single agents alone (> 50% growth inhibition). These are chlorambucil, tamoxifen citrate, fludarabine phosphate, exemestane, zoledronic acid, abiraterone, omacetaxine mepesuccinate, panobinostat, plerixafor, and acalabrutinib. All hit compounds, except omacetaxine mepesuccinate and panobinostat, had no effect on cell viability when applied alone with cell viabilities > 75% at 10 μM.

**Fig. 1:**
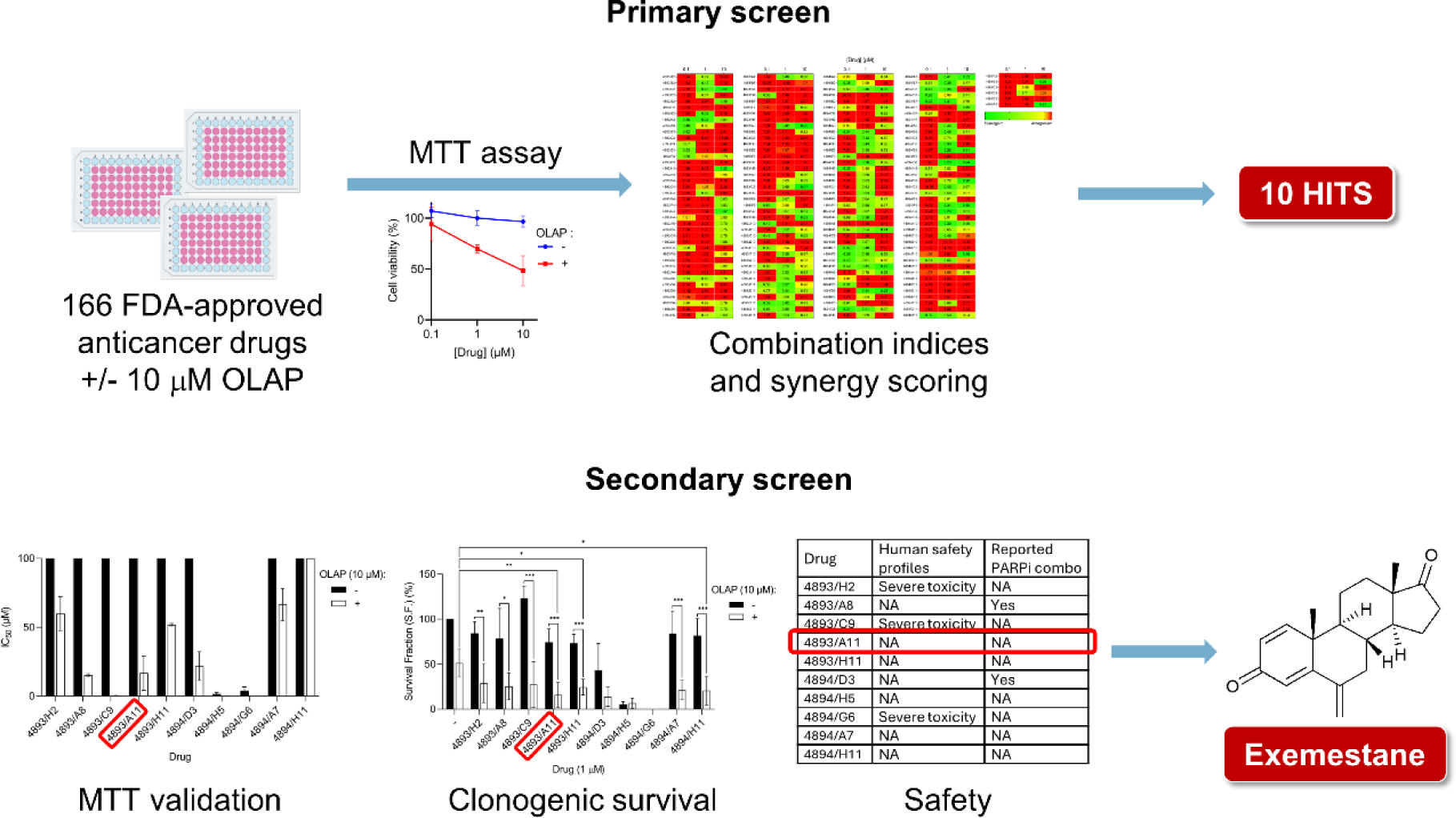
Schematic representation of the experimental steps implemented in the drug combination screening approach of Olaparib with a library of FDA-approved oncology drugs in MDA-MB-231 TNBC cells. The heatmap of combination indices (CIs), cell viability data, clonogenic survival assay data, and literature search results are shown in the supplementary information (Supplementary Table S1-S4 and Supplementary Fig. S1-S2).

To verify hits, these ten compounds were re-tested at a greater concentration range (0.01 to 100 μM) in the absence and presence of a low concentration (10 μM) of Olaparib for 24 h in MDA-MB-231 cells. Consistent with the initial findings, a substantial reduction in IC_50_ values (> 1.5- to > 200-fold change in IC_50_ values of combination vs. single agent) was observed following co-treatment with Olaparib compared to single agents alone, except for acalabrutinib, where IC_50_ concentrations of acalabrutinib alone or in combination are at > 100 μM (the maximum concentration tested; Supplementary Table S3). In addition to MTT assay, a long-term clonogenic survival assay was conducted. All hit compounds as single agents possess a low impact on colony formation of MDA-MB-231 cells where single agents Olaparib, exemestane, zoledronic acid and acalabrutinib gave S.F. of 51.4%, 74.3%, 73.2% and 81.7%, respectively (Supplementary Fig. S2). Strikingly, exemestane, zoledronic acid, and acalabrutinib showed a significant (*P* < 0.05) reduction in colony formation when in combination with Olaparib (S.F. of 15.9%, 24.6% and 20.7% for exemestane, zoledronic acid and acalabrutinib, respectively).

Finally, as the hit compounds are all FDA-approved drugs, a literature search was carried out to identify their drug classes, the human safety profiles, and whether prior PARPi combination studies of these compounds have been reported. Surprisingly, most of the hit compounds are pharmacologically diverse from each other (Supplementary Table S4). Moreover, studies have shown that chlorambucil, fludarabine phosphate, and panobinostat as monotherapies cause severe toxicity, limiting their use in clinics [22–24]. Interestingly, tamoxifen citrate (Clinicaltrials.gov identifier: NCT02093351) [25], and abiraterone (NCT03732820) [26], have been examined in combination with Olaparib in clinics. Therefore, these compounds were subsequently excluded. Collectively, Exemestane is identified as a new hit where no prior PARPi combinations have been reported with no reported severe toxicities as monotherapy. Moreover, significant decreased in cell survival verified the synergy observed between Olaparib and Exemestane, where a substantial reduction in IC_50_ values was observed (> 5.9-fold change; IC_50_ values = > 100 µM vs. 16.8 ± 12.5 µM for combination vs. single agent; *P* < 0.05; Fig. 2A and Supplementary Table S3). Besides, Olaparib and Exemestane combination treatment showed a significant decrease in clonogenicity (S.F. of 15.9% vs. 74.3% for combination vs. single agent; *P* < 0.01; Fig. 2B).

**Fig. 2:**
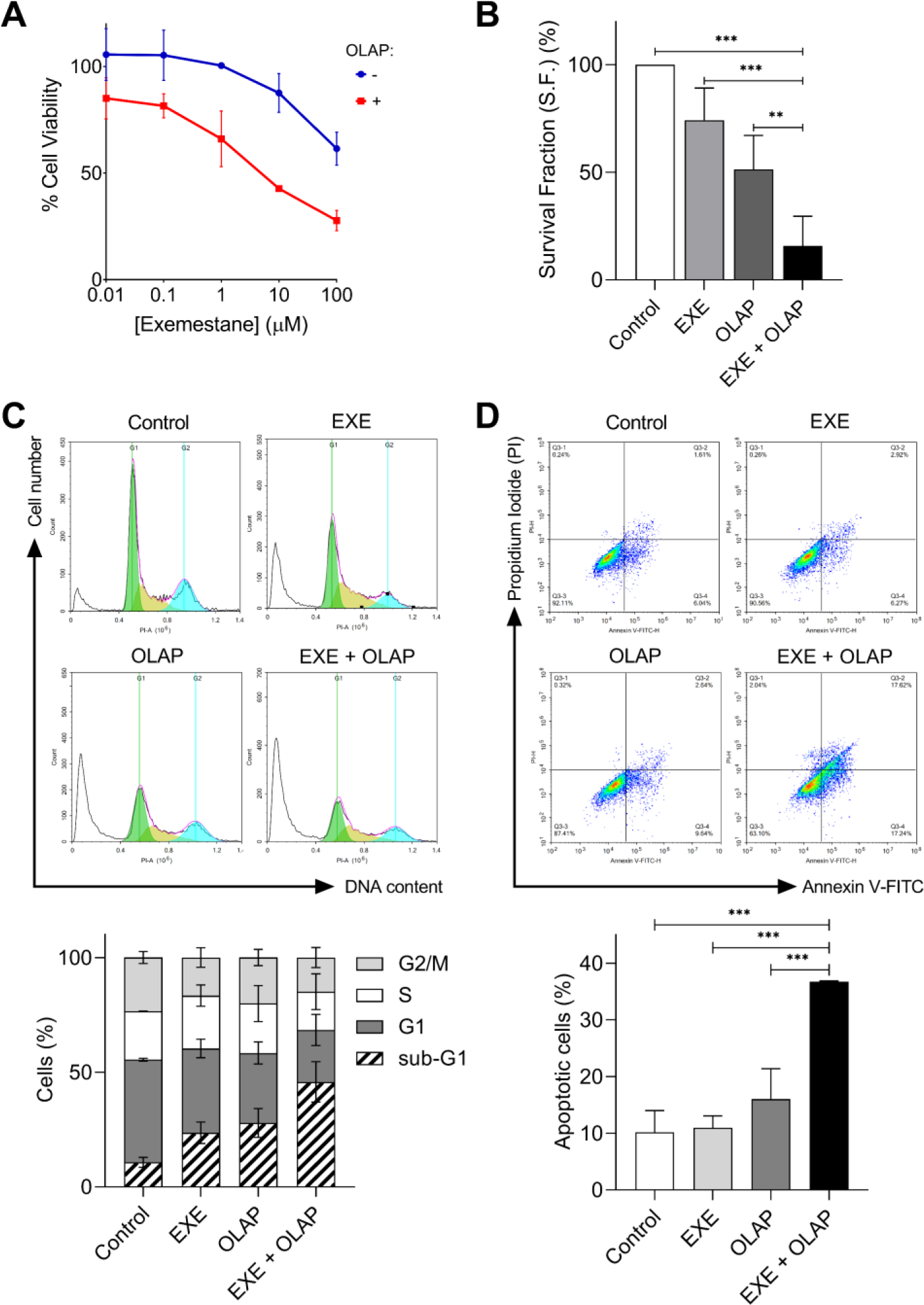
Exemestane synergized with Olaparib and leads to increased apoptosis in MDA-MB-231 cells. A) Cell viability of MDA-MB-231 cells following 24 h treatment with concentration gradients of Exemestane, with and without 10 µM Olaparib, as determined by MTT assay. B) Clonogenic survival assay of MDA-MB-231 cells treated with single-agent Exemestane in combination with Olaparib (24 h treatment). C) Cell cycle distribution of MDA-MB-231 cells treated with Exemestane (EXE, 25 µM) and Olaparib (OLAP, 10 µM), 72 h treatment, as determined by PI staining and flow cytometry. Upper, representative histograms, lower, quantification of cell cycle phase. D) Annexin V-FITC assay of MDA-MB-231 cells treated as in (C). Upper, representative scatterplots showing the percentage of cells in each quadrant, lower, quantification of apoptotic cells (Q3-2 and Q3-4 quadrants) for each treatment condition. Data expressed as mean ± SD of at three independent experiments. \*\*\**P* < 0.001 by ANOVA.

### Exemestane plus Olaparib leads to elevated apoptosis in MDA-MB-231 cells

To assess the cancer selectivity of the identified combination treatment, normal MCF10A breast epithelial cells were tested with Exemestane and Olaparib single agents or in combination. Normal MCF10A breast cells tolerated all combinations well, with cell viabilities > 70% for all concentrations tested (IC_50_s of > 100 µM for both single agent and combination; Supplementary Fig. S3). The minimal impact observed on normal breast epithelial cells indicates high cancer-selective activity of this treatment. Next, the cell cycle distribution following treatment was examined using flow cytometric analysis, where cells were treated with Exemestane and Olaparib alone or in combination for 72 h, at drug doses optimized for synergy. Notably, co-treatment with Exemestane and Olaparib caused a substantial increase in the proportion of sub-G1 phase cells (35.1% increase compared to untreated controls; Fig. 2C), indicative of elevated apoptosis. Quantifying apoptosis by Annexin V-FITC assay, single-agent treated groups showed comparable levels of apoptotic cells in comparison to untreated control (< 5.8% increase; *P* > 0.05; Fig. 2D), however, significantly higher levels of Annexin V-positive cells were apparent in the co-treated group, indicative of high levels of apoptosis (26.6% increase in comparison to untreated control; *P* < 0.001).

### Exemestane generates replication stress

Synergy with PARPi is commonly achieved by employing DNA-damaging agents that generate SSBs or replication fork arrest [27–29]. Accordingly, to explore the basis for the observed synergy, the roles of Exemestane in DDR activation was investigated. MDA-MB-231 cells were treated with a concentration gradient of Exemestane for 3 h and extracts were subjected to immunoblotting for several DDR signaling proteins, including activated (phosphorylated) p-ATR (at Thr1989) and p-ATM (at Ser1981), the two main upstream kinases in the signaling of DNA damage, and p-Chk1 (Ser345). Following short-term treatment with Exemestane, activation of the ATR signaling pathway was observed (1.9- to 2.3-fold increase in p-ATR/ATR for EXE vs. Control; *P* < 0.01; Fig. 3A). Basal phosphorylated (pATR) and endogenous levels of total ATR proteins were increased in correlation to replication stress [30]. It was also revealed that the p-Chk1 increased significantly (1.7- to 3.3-fold increase for EXE vs. Control; *P* < 0.01). These results indicate that Exemestane treatment leads to ATR/Chk1 signaling pathway activation, which responds to replication stress and increased DNA damage. Moreover, no activation of ATM was observed upon Exemestane treatment for 3 h where flow cytometric analysis showed that Exemestane-treated group had similar cell cycle profiles to untreated control (Supplementary Fig. S4). Strikingly, significant increase in total RPA1 foci formation upon Exemestane treatment was observed, where RPA1 is a single-stranded DNA (ssDNA) binding protein that stabilizes ssDNA during DNA damage repair and DNA replication (48.0% vs. 1.48% for EXE vs. Control; *P* < 0.01; Fig. 3B). Altogether, Exemestane was shown to target replication p-ATR causing replication defects without DSBs.

**Fig. 3:**
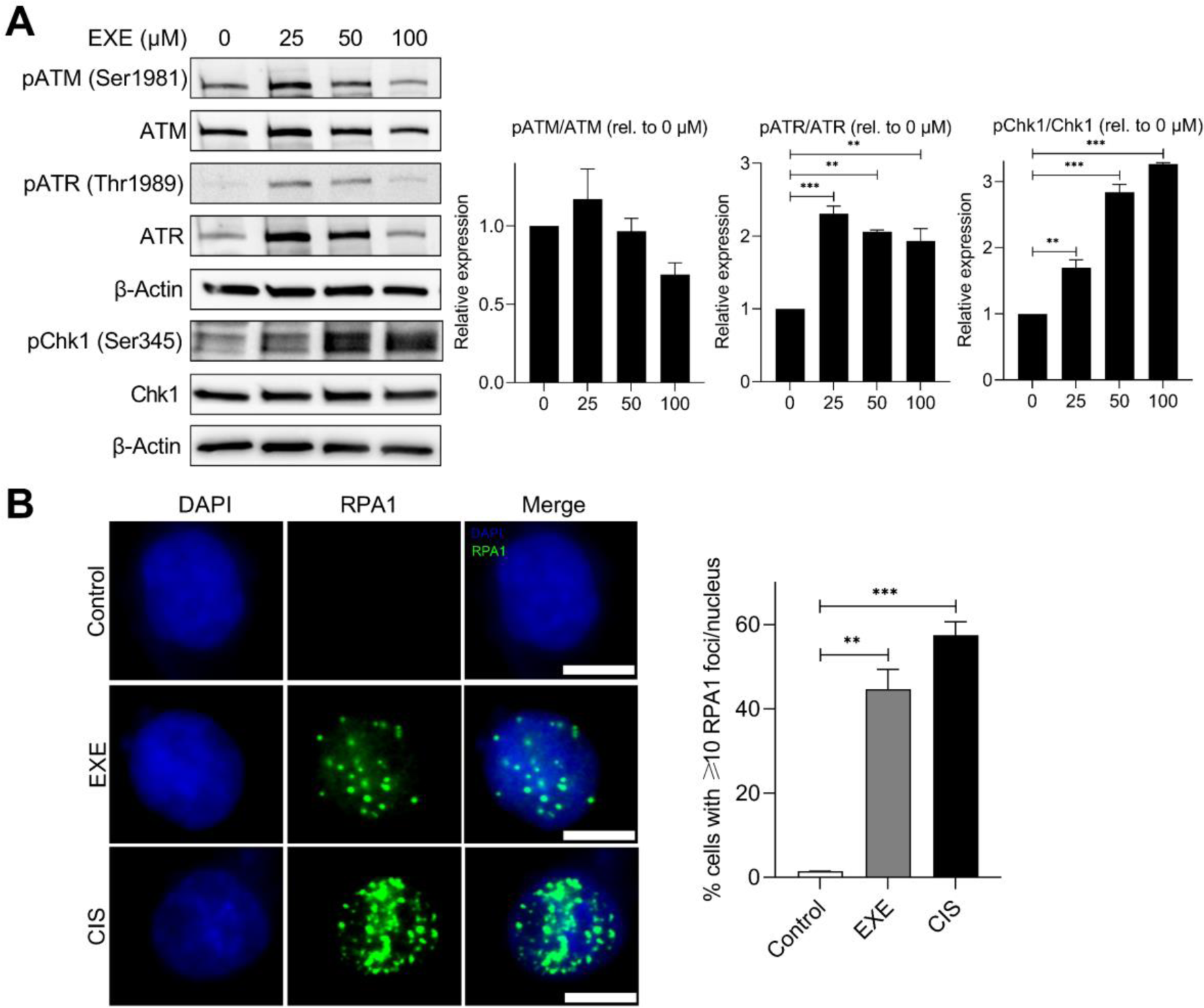
Exemestane single-agent generates replication stress. A) Western blot analysis of DDR activation following Exemestane treatment (25, 50 and 100 µM) for 3 h. Left, western blot images, right, quantification of protein bands from western blot images by densitometry. β-actin levels were monitored as a loading control. Uncropped blot images are shown in Supplementary Fig. S5. Data expressed as mean ± SD of duplicate experiments. B) RPA1 foci (green) formation in MDA-MB-231 cells nuclei following treatment with Exemestane (50 µM, 24 h). Cisplatin (CIS, 50 µM, 24 h) was employed as positive control. Nuclear staining by DAPI (blue) also included. Scale bars = 10 µm. Left, representative images, right, quantification of cells with more than 10 RPA1 foci where a minimum of 200 nuclei were counted for each independent experiment. Data expressed as mean ± SD of two independent experiments. \*\**P* < 0.01 and \*\*\**P* < 0.001 by ANOVA.

### Exemestane and Olaparib synergy accompanied by increased DNA DSB damage

We next evaluated the activation of the DDR signaling pathway after mono- and co-treatment with Exemestane and Olaparib. As expected, Exemestane- and Olaparib-treated groups showed a substantial increase in p-ATR/ATR in MDA-MB-231 cells (Fig. 4A). Notably, a significant (*P* < 0.05) reduction in p-ATR/ATR expression levels was observed upon co-treatment with Exemestane and Olaparib compared to single agents or untreated control groups, suggesting that the combination of these drugs decreases the Olaparib- or Exemestane-upregulation of p-ATR/ATR. Notably, the co-treatment of Exemestane and Olaparib showed a significant increase in γH2AX (at Ser139), an early DNA DSB marker levels, indicating increased DNA damage upon this combination treatment in MDA-MB-231 cells (2.6-fold increase for EXE + OLAP vs. NT; *P* < 0.001). In addition to immunoblotting, the presence of DSBs was examined by monitoring γH2AX levels via flow cytometry after 24 h of drug treatment, a time in which apoptosis is not yet observed. Consistent with the initial finding, Exemestane and Olaparib as single agents showed comparable levels of γH2AX compared to untreated control, indicating low DSB damage generated by these compounds at sub-cytotoxic concentrations (1.2-fold for EXE vs. Control; 1.7-fold for OLAP vs. NT; *P* > 0.05; Fig. 4B). Meanwhile, significantly greater γH2AX-positive cells are apparent upon combination treatment compared to their single-agent equivalents, indicating increased DNA DSB damage (5.8-fold for EXE + OLAP vs. Control; *P* < 0.01).

**Fig. 4:**
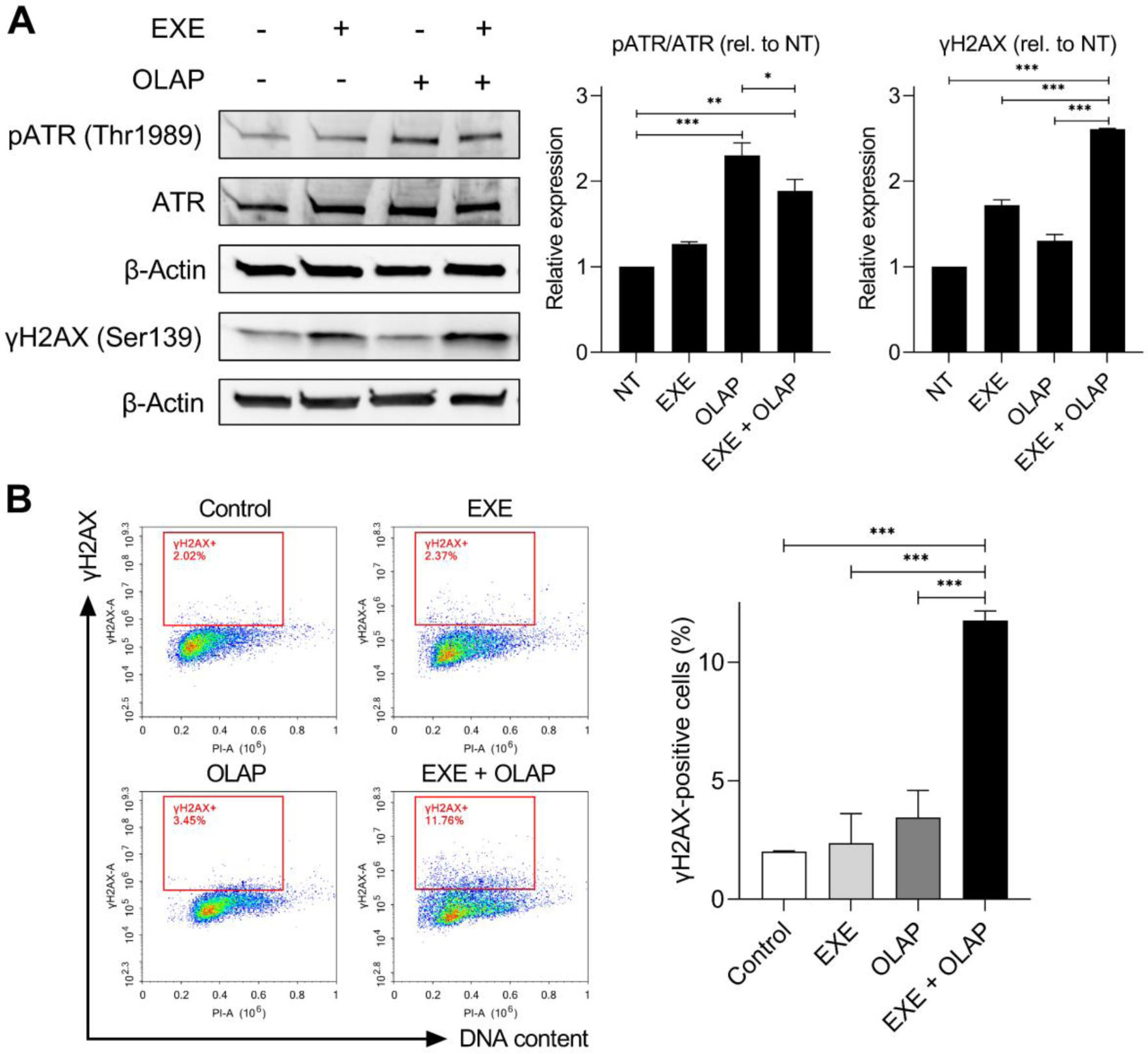
Exemestane and Olaparib synergy accompanied by increased DNA DSB damage. A) Western blot analysis of DDR activation following treatment with Exemestane (25 µM) and Olaparib (10 µM) for 3 h. Left, western blot images, right, quantification of protein bands from western blot images by densitometry. β-actin levels were monitored as a loading control. NT = untreated control. Uncropped blot images are shown in Supplementary Fig. S6. Data expressed as mean ± SD of duplicate experiments. B) γH2AX levels in cells treated as in (A) for 24 h, as determined by γH2AX/PI double staining. Left, the percentage of γH2AX-positive cells (red gating) in each population where y-axis shows the level of γH2AX immunofluorescence and x-axis shows DNA content (PI). Data expressed as mean ± SD of three independent experiments. \**P* < 0.05, \*\**P* < 0.01 and \*\*\**P* < 0.001 by ANOVA.

### Exemestane enhances the effects of Olaparib *in vivo*

To explore our findings *in vivo*, MDA-MB-231 xenograft tumor model of nude mice was established to investigate the synergistic effect of Exemestane and Olaparib on the growth of MDA-MB-231 cells *in vivo* (Fig. 5A). After 30 days of treatment, Exemestane (20 mg/kg) and Olaparib (50 mg/kg) treatment alone resulted in no significant difference in tumor volume and tumor weight compared to the vehicle-treated control mice (Fig. 5B-C). Strikingly, the combined treatment of Exemestane and Olaparib significantly (*P* < 0.05) inhibited tumor volume and tumor weight compared to single agents alone. Moreover, these findings were supported by the histologic analyses of the xenografts which showed a significant (*P* < 0.05) decrease in proliferation markers Ki67 and vascular endothelial marker CD31 along with an increase in γH2AX, the DNA damage marker, for the combination group (Fig. 5F). Further TUNEL staining analysis revealed significantly (*P* < 0.001) greater areas of apoptotic cells (green) in the combination group compared to single agents alone and control. Encouragingly, no loss of body weight of mice was observed following administrations of Exemestane, Olaparib, and their combination at the tested dosages (Fig. 5D). H&E-stained organs showed that these treatments did not cause significant damage to the heart, liver, spleen, lung, or kidney (Fig. 5E), suggesting the safety of this therapy.

**Fig. 5:**
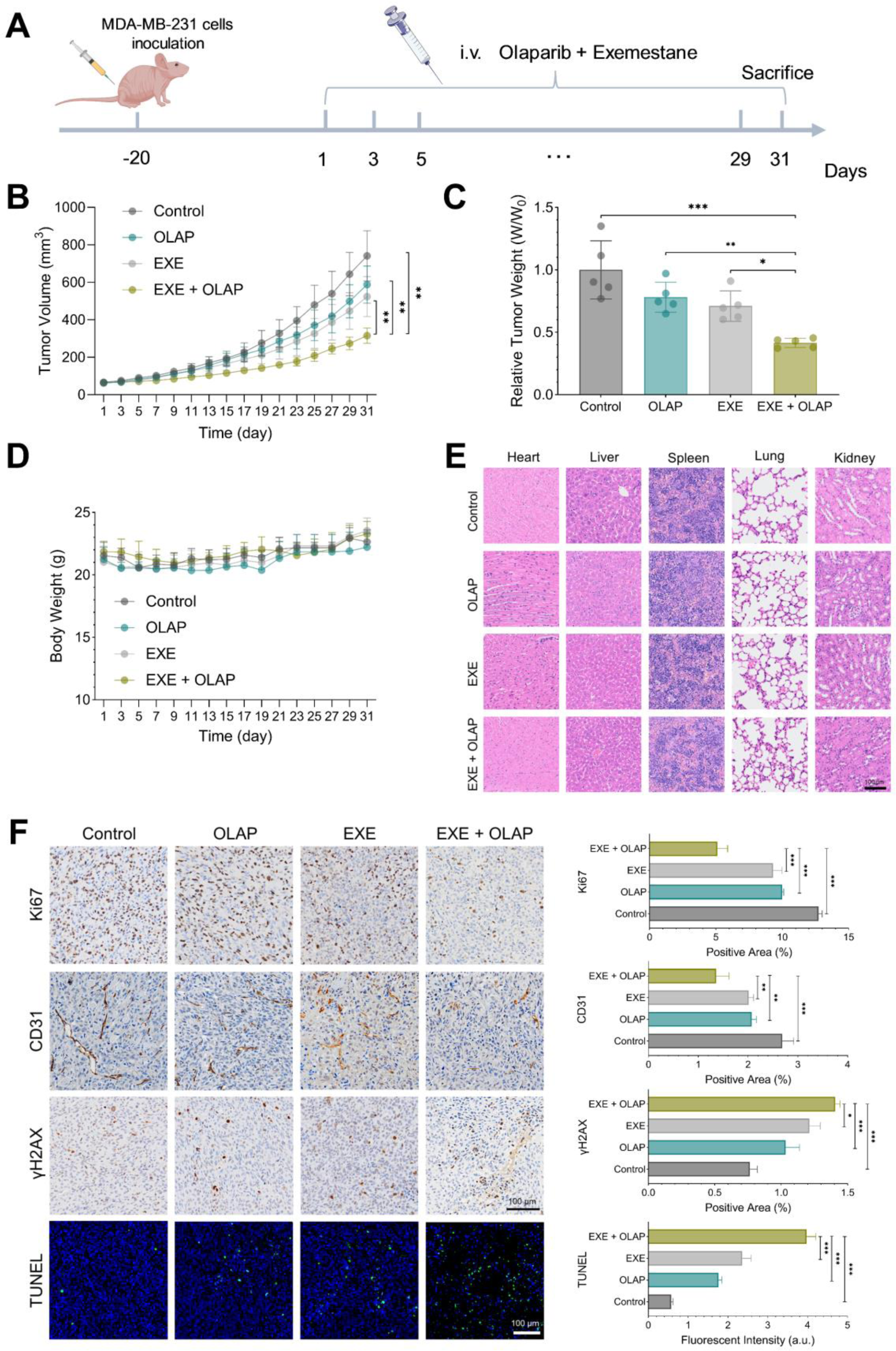
Exemestane enhances the effects of Olaparib *in vivo*. A) Schematic diagram indicating MDA-MB-231 cells were injected into nude mice and the mice were subsequently treated with Olaparib (50 mg/kg every two days for 30 days) and Exemestane (20 mg/kg every two days for 30 days) alone or in combination at the indicated times. B) Tumor volumes over 30 days of treatment with Olaparib and Exemestane alone or in combination (n = 5 per treatment group). C) Relative tumor weight of different treatment groups as indicated. D) Body weight of mice throughout the study. E) Representative images of H&E-staining of the tissue sections. Scale bars = 100 µm. F) Representative images of immunohistochemical staining for proliferation markers Ki67 (brown), vascular endothelial marker CD31 (brown) and DNA damage marker γH2AX, (brown) and TUNEL staining showing apoptotic cells (green). Nuclei staining (blue) also included. Left, representative images of xenografts used for evaluation, right, quantification of the percentage of positive area. Scale bars = 100 µm. Data are shown as the mean ± SD. \**P* < 0.05, \*\**P* < 0.01 and \*\*\**P* < 0.001 by ANOVA.

### Transcriptomic evaluations indicate alterations in pathways associated with cell death

To further explore the mechanism by which PARP inhibition combined with Exemestane shows enhanced efficacy in MDA-MB-231 tumors, transcriptome sequencing was conducted to analyze the differentially expressed genes (DEGs) between the treatment groups. A volcano plot illustrated changes in transcriptome expression, revealing 628 significantly upregulated and 907 downregulated genes in the Exemestane and Olaparib co-treated group compared to the control group (Fig. 6A). Since we previously showed that the co-treated group exhibited increased DNA damage, we screened DEGs associated with DNA repair to analyze their expression. Interestingly, the genes RAD21, MRE11, SIRT1 and Werner syndrome protein (WRN) were significantly (*P* < 0.05) upregulated, potentially indicating heightened DNA damage repair capacity in this group (Fig. 6B) [31, 32]. Subsequently, GO functional analysis and KEGG pathway enrichment analysis were conducted to compare the co-treated and control groups. GO analysis revealed that dysregulated DEGs were associated with pathways such as “regulation of cell cycle” and “regulation of intrinsic apoptosis signaling pathway in response to DNA damage” (Fig. 6C). Furthermore, KEGG analysis revealed enrichment of dysregulated DEGs in pathways including “phosphatidylinositol-3 kinase (PI3K)- Akt signaling pathways”, “nucleotide excision repair (NER)” and “P53 signaling pathways”, among others (Fig. 6D). Interestingly, the dysregulation of DEGs across all treatment groups reveals intriguing insights into the modulation of the PI3K-Akt signaling pathway (Supplementary Fig. S7), which plays a key role in cell survival and apoptosis [33, 34], and in drug resistance mechanisms [35, 36]. Olaparib treatment upregulates DEGs linked to the PI3K-Akt pathway, suggesting the activation of cell survival mechanisms. Conversely, Exemestane treatment downregulates DEGs associated with the PI3K-Akt pathway, potentially inhibiting this pro-survival signaling cascade. Strikingly, the co-treated group exhibited downregulation of DEGs in the PI3K-Akt pathway, indicating a synergistic effect in suppressing this pathway. Supporting this observation, a clustering heatmap analysis of DEGs associated with the PI3K-Akt signaling pathway revealed a significant (*P* < 0.05) increase in the expression of phosphatase and tensin homolog (PTEN) gene, a crucial tumor suppressor gene linked to negative regulation of PI3K-Akt pathway (Fig. 6E). In addition, the genes cyclin E1 (CCNE1), platelet derived growth factor receptor beta (PDGFRB), regulatory associated protein of MTOR complex 1 (RPTOR), and cyclin-dependent kinase 4 (CDK4) were found to be downregulated. These findings suggest that the combined inhibition by Exemestane and Olaparib disrupts key survival signals, leading to the suppression of the P13K-Akt pathway and thereby promoting cell death mechanisms via apoptosis in MDA-MB-231 tumors.

**Fig. 6:**
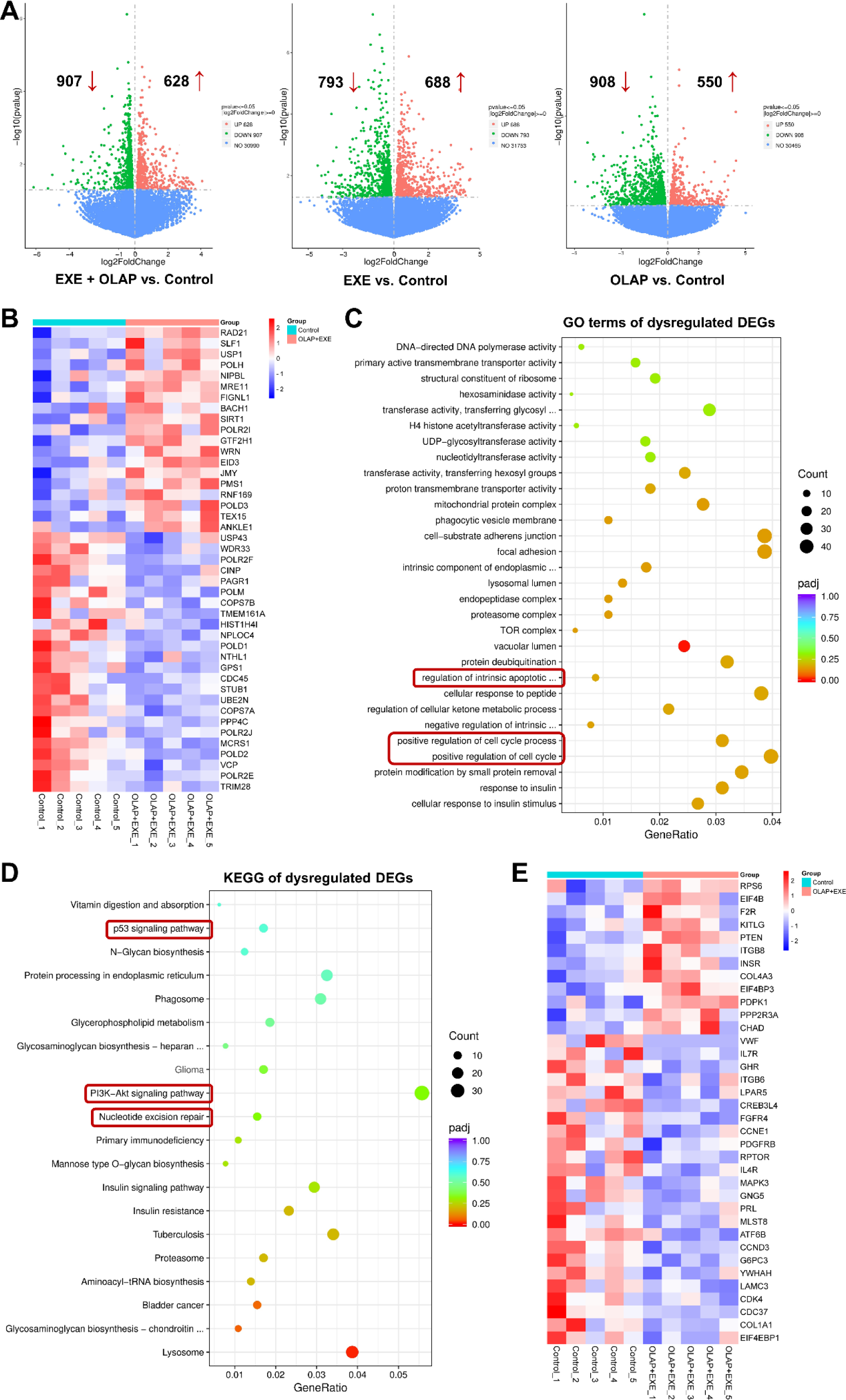
Transcriptomic evaluation following co-treatment with Exemestane and Olaparib. A) Volcano plot illustrating the differentially expressed genes (DEGs) across all treatment groups vs. control. Significantly upregulated (red) and downregulated (green) DEGs are shown. Log2 fold change (FC) = 0, n = 5 per group and *P* < 0.05 is considered statistically significant. B) Heatmap showing z-score normalized intensities of significantly upregulated and downregulated DEGs involved in DNA repair pathways for EXE+OLAP vs. control groups. *P* < 0.05 is considered statistically significant. C) GO enrichment analysis of DEGs across three functional categories: biological processes (BP), cellular components (CC), and molecular functions (MF). D) KEGG pathway enrichment analysis of DEGs in transcriptomes. The top 20 significantly enriched pathways are presented in order of enrichment score. Dot size represents gene count, and color represents the adjusted *P* value (Padj). E) Heatmap showing z-score normalized intensities of significantly upregulated and downregulated DEGs involved in PI3K-Akt signaling pathways for EXE+OLAP vs. control groups. *P* < 0.05 is considered statistically significant.

## DISCUSSION

The unbiased drug synergy screen employed in this study successfully isolated Exemestane, an aromatase inhibitor, as a synergistic pairing with Olaparib in BRCA-proficient MDA-MB-231 TNBC cells. Exemestane blocks the synthesis of estrogen and as monotherapy is an effective and selective treatment for postmenopausal patients with hormone-dependent breast cancers [37]. In patients with ER-positive cancers, Exemestane has also been examined in combination with other therapeutics including antimetabolites [38], alkylating agents [39], kinase inhibitors [40–42], and immune checkpoint antibodies [43, 44]. Although Exemestane has not been examined in combination with PARPi, a phase I study has examined Olaparib with other endocrine therapy drugs, including anastrozole, letrozole, or tamoxifen, in advanced solid tumor patients and in ER-positive breast cancer patients [25]. This combination was generally well tolerated, with no clinically relevant pharmacokinetic interactions identified. Our results both identify a new Exemestane drug combination and indicate the basis for the observed synergy is that Exemestane induces replication stress in BRCA-proficient cells - a previously unknown effect of the drug – and that the addition of a PARPi suppresses ATR activation, impairing DNA repair and resulting in the accumulation of cytotoxic DSBs, ultimately leading to apoptotic cell death. This serendipitous discovery into the polypharmacology of Exemestane validates our screening methodology.

The Exemestane-induced replication stress is a new discovery and the precise basis for this is unknown. It seems unlikely that Exemestane directly binds DNA and it is possible this occurs through the generation of reactive oxygen species (ROS), where in addition to blocking estrogen biosynthesis, Exemestane was found to generate ROS, resulting in the growth arrest of breast cancer [45]. Although only a single cancer cell line was examined in this study, it is encouraging that the combination showed no cytotoxicity in normal breast epithelial cells and that this combination inhibited the tumor growth of BRCA-proficient TNBC mouse xenograft model without noticeable signs of toxicity. The corresponding transcriptomic analysis provides further insights into the molecular mechanisms underlying the co-treatment of Exemestane and Olaparib in MDA-MB-231 tumors. Subsequent GO functional analysis and enriched KEGG pathways analysis of DEGs revealed dysregulation in cell survival and cell death pathways, further supporting our findings. Specifically, the enriched KEGG pathways of PI3K-Akt and P53 signaling influence apoptosis [33, 46], whereas nucleotide excision repair (NER) maintains genomic stability in response to DNA damage accumulation [47]. Interestingly, Olaparib treatment alone showed upregulation in the PI3K-Akt pathway, which is crucial for cell survival, consistent with prior studies [48, 49]. Other studies have also demonstrated that Olaparib treatment promotes the expression of genes involved in DNA repair pathways [50]. In contrast, KEGG analysis suggests that Exemestane indirectly downregulates the PI3K-Akt pathway in ER-negative breast cancer (TNBC), although this observation requires additional experimental confirmation. Notably, downregulation of PI3K-Akt pathway was observed in the co-treated group, indicating a synergistic effect in suppressing this pathway, possibly though p53 dysregulation [51]. These findings align with studies showing that inhibiting PI3K signaling (via BKM120) can sensitize BRCA-proficient TNBC cells to Olaparib [52, 53]. In addition, a phase I has examined the combination of Olaparib with the AKT inhibitor, capivasertib, in patients with and without BRCA mutations [54]. While further investigation is needed to fully understand the interaction between Exemestane and Olaparib in BRCA-proficient TNBC, these findings are consistent with our experimental results, which show extensive DNA damage beyond the repair capacity of the cells following co-treatment. Consequently, the cells activate programmed cell death mechanisms to prevent the propagation of damaged DNA, leading to enhanced apoptosis. While this provides a plausible explanation for the increased DNA damage and enhanced apoptosis observed in experiments following treatment with Exemestane and Olaparib in MDA-MB-231 tumors, further investigation across a greater range of cancer types is required.

In summary, this study supports the concept that a PARPi combination strategy represents a promising approach towards aggressive strains of TNBC, potentially extending the use of PARPi to BRCA-proficient cancers. Importantly, this newly identified synergistic combination may broaden the therapeutic benefits of Exemestane to include ER-negative breast cancer, thus potentially benefiting a broader population of breast cancer patients. However, optimal sequencing and combination strategies for Olaparib need to be further explored. Sequential therapy is often crucial in oncology, especially when combining different treatment modalities. However, determining the most effective sequence can be challenging and requires thorough investigation through extensive preclinical and clinical trials. For instance, the PARTNER clinical trial assessing neoadjuvant Olaparib with carboplatin/paclitaxel treatment in BRCA-proficient TNBC patients found no clinical benefit [55]. In another trial, carboplatin was administered prior to Olaparib in ovarian, breast and uterine cancer patients with and without BRCA mutation [56]. Strikingly, clinical activity was also seen in subsets of BRCA-proficient TNBC patients. These trials further underscore the necessity of refining the order in which treatments are given to maximize their effectiveness, as described in detail by Plana et al [57]. Nevertheless, the optimal sequence of Exemestane and Olaparib treatment is unknown, warranting additional investigation prior to clinical application.

## MATERIALS AND METHODS

### Materials and reagents

Antibodies for p-Chk1 (Ser345), p-ATR (Thr1989), total ATR, p-ATM (Ser 1981), total ATM, p-histone H2AX (Ser139), RPA1, β-actin, and HRP-linked secondary antibody were purchased from Cell Signaling Technology (CST). Antibodies for total Chk1, and Alexa-Fluor 488-conjugated secondary antibodies were purchased from Abcam. Olaparib and Exemestane were purchased from MedChemExpress (MCE). Cisplatin was purchased from SigmaAldrich. All other materials or reagents were purchased from SigmaAldrich and ThermoFisher Scientific, unless otherwise specified. Stock solutions of Olaparib and Exemestane were prepared at 10 mM in 100% dimethyl sulfoxide (DMSO) and were kept at -20 °C. Stock solutions of cisplatin (2 mM) were prepared in phosphate-buffered saline (PBS). 166 FDA-approved anticancer drugs were obtained from the Drug Synthesis and Chemistry Branch, Developmental Therapeutics Program, Division of Cancer Treatment and Diagnosis, National Cancer Institute (NCI) DTP Repository (Rockville, MD) as a plated set of compounds for research use. The details of each drug obtained from NCI DTP Repository, including their CAS number and molecular weight (MW), are described in Supplementary Table S1. The approved anticancer drugs set was provided in 96-well plates formatted with each well containing an individual compound at a volume of 20 μL and a concentration of 10 mM dissolved in 100% DMSO. These plated sets were kept at -20 °C. Compounds lacking long-term solubility in DMSO were suspended just before dispensing to avoid precipitation. Additionally, sub-stock solutions of the library were prepared at 0.1 mM and 1 mM to support low-concentration screening formats. All stock solutions were further diluted using PBS or Dulbecco’s modified Eagle’s medium (DMEM). The final DMSO concentration employed in the cell studies was < 0.1%.

### Cell lines and culture conditions

MDA-MB-231 TNBC cells were cultured in DMEM supplemented with 10% fetal bovine serum (FBS) and 1% penicillin/streptomycin (P/S) antibiotic. The MCF10A normal breast cell line was cultured in DMEM supplemented with 5% horse serum, 0.5 μg/mL hydrocortisone, 20 ng/mL recombinant human EGF (hEGF), 10 μg/mL insulin, and 1% P/S antibiotic. Cells were cultured as monolayers and maintained in a cell culture incubator at 37 °C under a humidified atmosphere containing 5% carbon dioxide (CO_2_). They were routinely subcultured with trypsin approximately twice a week at 80-90% confluency. Unless stated otherwise, all treatments were started 24 h after seeding.

### MTT assay

MDA-MB-231 TNBC and MCF10A normal cells were seeded (1 x 10^4^ cells/well) in 96-well plates and were treated as stated in the main text. For the drug combination screen, the working solution of each drug treatment at 10 µM and the subsequent 10-fold dilutions were prepared. Single drugs as well as combinations were administered for 24 h. Each experimental 96-well plate contained drug treatments, each in triplicate. The negative control (untreated cells, 0% inhibition) and positive control (Triton X-100, 100% inhibition or complete cell death) were included in each plate. Following treatment at selected time points, solutions were removed, thiazolyl blue tetrazolium bromide (MTT, 0.5 mg/mL) reagent was added to the cells, and plates were incubated for 3 h. The reduced purple formazan crystals were solubilized with 100 μL of DMSO, and the absorbance at 570 nm (620 nm as the reference wavelength) was measured using a microplate reader (Tecan Infinite F50). The half-maximal inhibitory concentration (IC_50_) for inhibiting cell viability for each compound was derived from the dose-response curves using GraphPad Prism version 9.00 for Windows, GraphPad Software, La Jolla, California, USA, www.graphpad.com.

### Drug interaction analysis

For drug interaction analysis, dose-effect curves for mono- and co-treatments were generated from MTT assay data, and the combination index (CI) values for each combination were calculated using CalcuSyn and CompuSyn software (Biosoft, Cambridge, UK) as established by Chou and Talalay [20, 21]. CI < 0.9 indicates synergism, CI = 0.9-1.0 indicates additive, and CI > 1 indicates antagonism. GraphPad Prism Software was used to generate a three-color scale based on CI values obtained, where synergism is represented by green, additive by yellow, and antagonism by red. The colors of each CI value were interpolated between these constraints accordingly. Ten hit compounds were isolated from the initial primary screen for further validation.

### Clonogenic survival assay

MDA-MB-231 cells were seeded (1 x 10^3^ cells/well) in 6-well plates and were treated as stated in the main text. After treatment, solutions were removed, and cells were cultured in a compound-free medium for 10-14 days to allow colony formation. Cells were then washed twice with 1X PBS, fixed with ice-cold 100% methanol for 15-20 min at 4°C, and stained with a 0.5% crystal violet solution for 20 min. The staining solution was washed with water, and images were photographed with a digital camera. Individual colonies were counted using ImageJ software, and the survival fraction was determined (normalized to controls).

### Cell cycle analysis

MDA-MB-231 cells were seeded (3 x 10^5^ cells/well) in 6-well plates and were treated as stated in the main text. Following treatment, cells were trypsinized and washed with 1X PBS twice. This was followed by fixation in ice-cold 70% ethanol for at least overnight at 4 °C. After fixation, fixed cells were centrifuged (1,000 rpm, 5 min), and the resulting cell pellets were washed with 1X PBS twice. Samples were resuspended in 500 µL of 1X PBS containing 100 µg/ml RNase A solution for 15 min. Samples were then stained with propidium iodide (PI) (20 µg/mL, in the dark) at 37 °C. After incubation, samples were stored on ice until data acquisition by flow cytometry using a NovoCyte flow cytometer (Agilent Technologies, USA), and NovoExpress software. For each sample, a minimum of 10,000 cells was counted.

### Apoptosis Annexin V-FITC/PI assay

MDA-MB-231 cells were seeded (3 x 10^5^ cells/well) in 6-well plates and were treated as stated in the main text. After treatment, cells were trypsinized and washed with 1X PBS twice. This was followed by the addition of 500 µL 1X binding buffer and 5 µL Annexin V-FITC (Elabscience). The cell-containing mixture was incubated for 20 min at RT. 5 µL of PI (20 µg/mL) was added before flow cytometric analysis using a flow cytometer, and results were analyzed using NovoExpress software. For each sample, a minimum of 10,000 cells was counted.

### Immunoblotting

MDA-MB-231 cells were seeded (8 x 10^5^ cells/dish) in 60 mm cell culture dish, allowed to adhere for 24 h, and were treated as stated in the main text. Following treatment, cells were harvested and lysed in 1X RIPA (radioimmunoprecipitation assay) buffer (CST) supplemented with protease inhibitors and phosphatase inhibitors. Aliquots of cell lysates (40 µg total protein) were separated by sodium dodecyl sulfate-polyacrylamide gel electrophoresis (SDS-PAGE) using 4-20% Mini-PROTEAN TGX precast protein gels (Bio-Rad) and were transferred onto a nitrocellulose membrane using wet western blot transfer. Following this, the membrane was blocked in blocking buffer (5% BSA in TBS-T (0.1% Tween 20 in 1X TBS)) for 1 h at RT. Membranes were then washed once with TBS-T (5 min) and probed with primary antibodies in 5% BSA in TBS-T solutions, overnight, at 4 °C. The primary antibodies used were: p-Chk1 (Ser345) (1:500), total Chk1 (1:1,000), p-ATR (Thr1989) (1:500), total ATR (1:1,000), p-ATM (Ser 1981) (1:500), total ATM (1:1,000), p-histone H2AX (Ser139) (1:1,000), and β-actin (1:1,000). Membranes were then washed with TBS-T (3 x 5 min) and probed with a suitable horseradish peroxidase (HRP)-conjugated secondary antibody in 5% BSA in TBS-T solutions (1:3,000) for 1 h at RT. Membranes were then washed (TBS-T, 3 x 5 min) and were incubated with chemiluminescence substrates of SignalFire ECL reagent (CST) or WesternBright ECL HRP substrate (Advansta) for 1 min at RT. Protein expression was visualized using a Syngene G:Box gel documentation system. ImageJ software was used for densitometry data acquisition. β-actin was used as a loading control.

### Immunofluorescence

Cells were seeded in a 60 mm cell culture dish on 24 x 24 mm coverslips at a seeding density of 8 x 10^5^ cells/dish and were treated as stated in the main text. Solution was removed, cells were washed with ice-cold 1X PBS twice, and fixed with 4% paraformaldehyde (PFA, 15 min, RT). After fixation, PFA was removed and cells were washed with ice-cold 1X PBS twice. Cells were then permeabilized with Triton X-100 (0.5% in PBS, 15 min, on ice) and washed with ice-cold 1X PBS thrice. Samples were blocked with 3% BSA in PBS-T (PBS with 0.1% Tween 20) for 1 h before incubation with the primary antibody (anti-RPA1, 1:50) diluted in 3% BSA in PBS-T overnight, in a humid chamber at 4 °C. Following incubation, samples were washed in PBS-T (3 x 5 min) and incubated with Alexa Fluor 488-conjugated secondary antibodies (1/200 dilution in 3% BSA in PBS-T) for 1 h, protected from light at RT. After further washing with PBS-T (3 x 5 min), samples were co-stained with DAPI (5 µg/mL, 2 min). Coverslips were mounted onto glass slides with antifade mounting medium and cells were visualized by confocal microscopy in a dark room. Microscopy images were processed, and the number of cells with more than 10 RPA1 foci were counted using ImageJ software. A minimum of 200 nuclei were counted for each independent experiment.

### Flow cytometry measurement for γH2AX

MDA-MB-231 cells were seeded at 3 x 10^5^ cells/well in 6-well plates and were treated as stated in the main text. Following treatment, cells were trypsinized and washed with 1X PBS twice. Cells were then fixed with 4% PFA for 15 min at RT. Following fixation, cells were washed with 1X PBS and resuspended in 100 µL of 1X PBS. Thereafter, cells were permeabilized by adding ice-cold 100% methanol slowly to pre-chilled cells, while gently vortexing to a final concentration of 90% methanol and left at -20°C for 2 h. Cells were then washed in excess 1X PBS and incubated with diluted primary antibody (γH2AX, 1:400) in antibody dilution buffer (0.5% BSA in PBS) overnight at 4 °C. Following incubation, cells were washed with antibody dilution buffer and incubated with diluted fluorochrome-conjugated secondary antibody for 1 h at RT, in the dark (Alexa-Fluor 488, 1:2,000). Next, samples were washed with antibody dilution buffer and resuspended in 500 µL 1X PBS containing RNase A solution (10 μg/mL, 15 min, RT). Samples were then stained with PI (20 μg/mL, 30 min, in the dark). Samples were acquired and analyzed with a NovoCyte flow cytometer and NovoExpress software. For each sample, a minimum of 10,000 cells was counted.

### *In vivo* xenograft model

5 x 10^6^ MDA-MB-231 cells were injected subcutaneously into the right hind limb of female nude mice to establish the tumor model (n=5). Upon tumor formation after 20 days and when the tumor volume reaches about 70 mm^3^, the mice were randomly divided into four following groups (Control, Olaparib, Exemestane and their combination, n=5 per treatment group). Olaparib was administered at 50 mg/kg and Exemestane was administered at 20 mg/kg, every 2 days for 30 days by intraperitoneal injection. Physiological saline was used as a solvent control. Tumor growth was assessed every two days for the following 30 days using vernier calipers, and tumor volume was calculated using the formula: tumor volume = (length × width^2^)/2. The body weight of the mice was recorded every two days for a total of 30 days. At the end of the 30-days period, the mice were euthanized, one day after the last treatment, and the heart, liver, spleen, lungs, kidneys, and tumors were collected for further analysis. In this study, animal experiments were conducted in accordance with the protocol approved by the Animal Ethics Committee of West China Hospital, Sichuan University (Approval No. 20230726001).

### Histological and immunohistochemical (IHC) staining analysis

Tissue samples were fixed in 4% paraformaldehyde and embedded in paraffin. Paraffin blocks were sectioned into 5–6 μm thickness, placed onto slides and stained with haematoxylin and eosin (H&E, servicebio). For immunohistochemical staining analysis, slides were incubated with primary antibodies against Ki-67 (servicebio, GB111499, 1:1,000), CD31 (servicebio, GB113151, 1:200) and γH2AX (servicebio, GB111841, 1:100) overnight at 4 °C. Slides were then incubated with the anti-rabbit HRP secondary antibody (servicebio, GB23303, 1:200) of the corresponding species of the primary antibody for 50 minutes, and developed using 3,3′- diaminobenzidine (DAB, servicebio). The slides of H&E and immunohistochemical staining assays were imaged using the SLIDEVIEW VS200 slide scanner (Olympus). Apoptosis was detected using a one-step TUNEL apoptosis assay kit according to the manufacturer’s instructions (Wuhan Servicebio Technology Co. Ltd., Hubei, China), and samples were observed under a Leica confocal microscopy (Stellaris 8). Nuclei were counterstained with haematoxylin or DAPI for IHC and the TUNEL reaction, respectively. Three images were randomly captured per slide and the percentage of stained area was analysed by ImageJ software.

### Transcriptomics analysis

Briefly, tissue samples were subjected to LC-MS/MS analysis for omics evaluation. Transcriptome analyses was performed using the omics technology platform at Novogene Bioinformatics Technology Co., Ltd. (Beijing, China). Differentially expressed genes (DEGs) were analyzed using volcano plot analysis, clustered heat map analysis, Gene Ontology (GO) functional analysis, and KEGG (Kyoto Encyclopedia of Genes and Genomes) pathway enrichment analysis, all conducted with NovoMagic software and https://www.bioinformatics.com.cn, an online platform for data analysis and visualization.

### Statistical analysis

Unless stated otherwise, the representation of figures and all data were statistically analyzed using GraphPad Prism software. All experiments were performed in triplicate and repeated three independent times (n=3), unless specified otherwise. Each data point represents the mean value ± standard deviation (SD). Statistical significance between the groups was analyzed using one-way analysis of variance (ANOVA). Differences were considered significant when the *P* values were less than 0.05.

## DATA AVAILABILITY

Any data generated from these studies is available from the corresponding author upon reasonable request.

## ACKNOWLEDGEMENTS

This work was supported by the Welsh Government and a Sêr Cymru Strategic Partner Acceleration Award (80761-SU-242) and Universiti Putra Malaysia through Geran Putra Inisiatif Siswazah (GP-IPS/2022/9737200). The PI wishes to thank the National Cancer Institute Developmental Therapeutics Program (NCI/DTP) https://dtp.cancer.gov for providing the FDA-approved anticancer drugs set used in this manuscript.

## AUTHOR CONTRIBUTIONS

MG, HA, and XT conceived and designed the project. NAY and LS performed the experiments, data curation, data analysis, and data interpretation. Resources and supervision were provided by MG, HA, XT, and SLC. NAY and MG wrote the original draft of the manuscript. All authors revised and approved the final version of the published manuscript.

## REFERENCES

1. Zagami P, Carey LA. Triple negative breast cancer: Pitfalls and progress. npj Breast Cancer. 2022;8:95.

2. Yin L, Duan J-J, Bian X-W, Yu S-c. Triple-negative breast cancer molecular subtyping and treatment progress. Breast Cancer Res. 2020;22:61.

3. Bianchini G, De Angelis C, Licata L, Gianni L. Treatment landscape of triple-negative breast cancer — expanded options, evolving needs. Nat Rev Clin Oncol. 2022;19:91–113.

4. Mateo J, Lord CJ, Serra V, Tutt A, Balmaña J, Castroviejo-Bermejo M, et al. A decade of clinical development of PARP inhibitors in perspective. Ann Oncol. 2019;30:1437–1447.

5. Lord CJ, Ashworth A. PARP inhibitors: Synthetic lethality in the clinic. Science. 2017;355:1152–1158.

6. Curtin NJ, Szabo C. Poly(ADP-ribose) polymerase inhibition: past, present and future. Nat Rev Drug Discov. 2020;19:711–736.

7. Gonzalez-Angulo AM, Timms KM, Liu S, Chen H, Litton JK, Potter J, et al. Incidence and outcome of BRCA mutations in unselected patients with triple receptor-negative breast cancer. Clin Cancer Res. 2011;17:1082–1089.

8. Engel C, Rhiem K, Hahnen E, Loibl S, Weber KE, Seiler S, et al. Prevalence of pathogenic BRCA1/2 germline mutations among 802 women with unilateral triple-negative breast cancer without family cancer history. BMC Cancer. 2018;18:265.

9. Lopez JS, Banerji U. Combine and conquer: challenges for targeted therapy combinations in early phase trials. Nat Rev Clin Oncol. 2017;14:57–66.

10. Jaaks P, Coker EA, Vis DJ, Edwards O, Carpenter EF, Leto SM, et al. Effective drug combinations in breast, colon and pancreatic cancer cells. Nature. 2022;603:166–173.

11. Bhamidipati D, Haro-Silerio JI, Yap TA, Ngoi N. PARP inhibitors: enhancing efficacy through rational combinations. Br J Cancer. 2023;129:905–916.

12. Prasad CB, Prasad SB, Yadav SS, Pandey LK, Singh S, Pradhan S, et al. Olaparib modulates DNA repair efficiency, sensitizes cervical cancer cells to cisplatin and exhibits anti-metastatic property. Sci Rep. 2017;7:12876.

13. Shen YT, Evans JC, Zafarana G, Allen C, Piquette-Miller M. BRCA Status Does Not Predict Synergism of a Carboplatin and Olaparib Combination in High-Grade Serous Ovarian Cancer Cell Lines. Mol Pharm. 2018;15:2742–2753.

14. Eetezadi S, Evans JC, Shen YT, De Souza R, Piquette-Miller M, Allen C. Ratio-Dependent Synergism of a Doxorubicin and Olaparib Combination in 2D and Spheroid Models of Ovarian Cancer. Mol Pharm. 2018;15:472–485.

15. Jiang Y, Dai H, Li Y, Yin J, Guo S, Lin SY, et al. PARP inhibitors synergize with gemcitabine by potentiating DNA damage in non-small-cell lung cancer. Int J Cancer. 2019;144:1092–1103.

16. Kim H, Xu H, George E, Hallberg D, Kumar S, Jagannathan V, et al. Combining PARP with ATR inhibition overcomes PARP inhibitor and platinum resistance in ovarian cancer models. Nat Commun. 2020;11:3726.

17. Schmidt L, Kling T, Monsefi N, Olsson M, Hansson C, Baskaran S, et al. Comparative drug pair screening across multiple glioblastoma cell lines reveals novel drug-drug interactions. Neuro Oncol. 2013;15:1469–1478.

18. Ariey-Bonnet J, Berges R, Montero M-P, Mouysset B, Piris P, Muller K, et al. Combination drug screen targeting glioblastoma core vulnerabilities reveals pharmacological synergisms. EBioMedicine. 2023;95.

19. Lui GYL, Shaw R, Schaub FX, Stork IN, Gurley KE, Bridgwater C, et al. BET, SRC, and BCL2 family inhibitors are synergistic drug combinations with PARP inhibitors in ovarian cancer. EBioMedicine. 2020;60:102988.

20. Chou T-C, Talalay P. Analysis of combined drug effects: a new look at a very old problem. Trends Pharmacol Sci. 1983;4:450–454.

21. Chou T-C, Talalay P. Quantitative analysis of dose-effect relationships: the combined effects of multiple drugs or enzyme inhibitors. Adv Enzyme Regul. 1984;22:27–55.

22. Palmer RG, Denman AM. Malignancies induced by chlorambucil. Cancer Treat Rev. 1984;11:121–129.

23. Chun HG, Leyland-Jones BR, Caryk SM, Hoth DF. Central nervous system toxicity of fludarabine phosphate. Cancer Treat Rep. 1986;70:1225–1228.

24. Eleutherakis-Papaiakovou E, Kanellias N, Kastritis E, Gavriatopoulou M, Terpos E, Dimopoulos MA. Efficacy of Panobinostat for the Treatment of Multiple Myeloma. J Oncol. 2020;2020:7131802.

25. Plummer R, Verheul HM, De Vos F, Leunen K, Molife LR, Rolfo C, et al. Pharmacokinetic Effects and Safety of Olaparib Administered with Endocrine Therapy: A Phase I Study in Patients with Advanced Solid Tumours. Adv Ther. 2018;35:1945–1964.

26. Clarke NW, Armstrong AJ, Thiery-Vuillemin A, Oya M, Shore N, Loredo E, et al. Abiraterone and Olaparib for Metastatic Castration-Resistant Prostate Cancer. NEJM Evidence. 2022;1:EVIDoa2200043.

27. Yusoh NA, Tiley PR, James SD, Harun SN, Thomas JA, Saad N, et al. Discovery of Ruthenium(II) Metallocompound and Olaparib Synergy for Cancer Combination Therapy. J Med Chem. 2023;66:6922–6937.

28. Yusoh NA, Leong SW, Chia SL, Harun SN, Rahman MBA, Vallis KA, et al. Metallointercalator [Ru(dppz)2(PIP)]2+ Renders BRCA Wild-Type Triple-Negative Breast Cancer Cells Hypersensitive to PARP Inhibition. ACS Chem Biol. 2020;15:378–387.

29. Yusoh NA, Chia SL, Saad N, Ahmad H, Gill MR. Synergy of ruthenium metallo-intercalator, [Ru(dppz)2(PIP)]2+, with PARP inhibitor Olaparib in non-small cell lung cancer cells. Sci Rep. 2023;13:1456.

30. Igarashi T, Mazevet M, Yasuhara T, Yano K, Mochizuki A, Nishino M, et al. An ATR-PrimPol pathway confers tolerance to oncogenic KRAS-induced and heterochromatin-associated replication stress. Nat Commun. 2023;14:4991.

31. D’Amours D, Jackson SP. The MRE11 complex: at the crossroads of DNA repair and checkpoint signalling. Nat Rev Mol Cell Biol. 2002;3:317–327.

32. Alves-Fernandes DK, Jasiulionis MG. The Role of SIRT1 on DNA Damage Response and Epigenetic Alterations in Cancer. Int J Mol Sci. 2019;20:3153.

33. He Y, Sun MM, Zhang GG, Yang J, Chen KS, Xu WW, et al. Targeting PI3K/Akt signal transduction for cancer therapy. Signal Transduct Target Ther. 2021;6:425.

34. Franke TF, Hornik CP, Segev L, Shostak GA, Sugimoto C. PI3K/Akt and apoptosis: size matters. Oncogene. 2003;22:8983–8998.

35. Dong C, Wu J, Chen Y, Nie J, Chen C. Activation of PI3K/AKT/mTOR Pathway Causes Drug Resistance in Breast Cancer. Front Pharmacol. 2021;12:628690.

36. Liu R, Chen Y, Liu G, Li C, Song Y, Cao Z, et al. PI3K/AKT pathway as a key link modulates the multidrug resistance of cancers. Cell Death Dis. 2020;11:797.

37. Buzdar AU, Robertson JF, Eiermann W, Nabholtz JM. An overview of the pharmacology and pharmacokinetics of the newer generation aromatase inhibitors anastrozole, letrozole, and exemestane. Cancer. 2002;95:2006–2016.

38. Alvarado-Miranda A, Lara-Medina FU, Muñoz-Montaño WR, Zinser-Sierra JW, Galeana PAC, Garza CV, et al. Capecitabine Plus Aromatase Inhibitor as First Line Therapy for Hormone Receptor Positive, HER2 Negative Metastatic Breast Cancer. Curr Oncol. 2023;30:6097–6110.

39. Young PA, Márquez-Garbán DC, Noor ZS, Moatamed N, Elashoff D, Grogan T, et al. Investigation of Combination Treatment With an Aromatase Inhibitor Exemestane and Carboplatin-Based Therapy for Postmenopausal Women With Advanced NSCLC. JTO Clin Res Rep. 2021;2:100150.

40. Baselga J, Campone M, Piccart M, Burris HA, 3rd, Rugo HS, Sahmoud T, et al. Everolimus in postmenopausal hormone-receptor-positive advanced breast cancer. N Engl J Med. 2012;366:520–529.

41. Jerusalem G, de Boer RH, Hurvitz S, Yardley DA, Kovalenko E, Ejlertsen B, et al. Everolimus Plus Exemestane vs Everolimus or Capecitabine Monotherapy for Estrogen Receptor-Positive, HER2-Negative Advanced Breast Cancer: The BOLERO-6 Randomized Clinical Trial. JAMA Oncol. 2018;4:1367–1374.

42. Bardia A, Hurvitz SA, DeMichele A, Clark AS, Zelnak A, Yardley DA, et al. Phase I/II Trial of Exemestane, Ribociclib, and Everolimus in Women with HR(+)/HER2(-) Advanced Breast Cancer after Progression on CDK4/6 Inhibitors (TRINITI-1). Clin Cancer Res. 2021;27:4177–4185.

43. Chen IC, Lin C-H, Chang D-Y, Chen TW-W, Wang M-Y, Ma W-L, et al. Abstract CT028: A pilot study of pembrolizumab and exemestane/leuprolide in premenopausal hormone receptor positive/HER2 negative locally advanced or metastatic breast cancer (PEER). Cancer Res. 2021;81:CT028–CT028.

44. Ge X, Yost SE, Lee JS, Frankel PH, Ruel C, Cui Y, et al. Phase II Study Combining Pembrolizumab with Aromatase Inhibitor in Patients with Metastatic Hormone Receptor Positive Breast Cancer. Cancers (Basel). 2022;14:4279.

45. Nuvoli B, Camera E, Mastrofrancesco A, Briganti S, Galati R. Modulation of reactive oxygen species via ERK and STAT3 dependent signalling are involved in the response of mesothelioma cells to exemestane. Free Radic Biol Med. 2018;115:266–277.

46. Aubrey BJ, Kelly GL, Janic A, Herold MJ, Strasser A. How does p53 induce apoptosis and how does this relate to p53-mediated tumour suppression? Cell Death Differ. 2018;25:104–113.

47. Marteijn JA, Lans H, Vermeulen W, Hoeijmakers JH. Understanding nucleotide excision repair and its roles in cancer and ageing. Nat Rev Mol Cell Biol. 2014;15:465–481.

48. Jagtap P, Szabó C. Poly(ADP-ribose) polymerase and the therapeutic effects of its inhibitors. Nat Rev Drug Discov. 2005;4:421–440.

49. Gallyas F, Jr., Sumegi B, Szabo C. Role of Akt Activation in PARP Inhibitor Resistance in Cancer. Cancers (Basel). 2020;12:532.

50. Wang S-P, Wu S-Q, Huang S-H, Tang Y-X, Meng L-Q, Liu F, et al. FDI-6 inhibits the expression and function of FOXM1 to sensitize BRCA-proficient triple-negative breast cancer cells to Olaparib by regulating cell cycle progression and DNA damage repair. Cell Death Dis. 2021;12:1138.

51. Bozulic L, Surucu B, Hynx D, Hemmings BA. PKBα/Akt1 Acts Downstream of DNA-PK in the DNA Double-Strand Break Response and Promotes Survival. Mol Cell. 2008;30:203–213.

52. Li Y, Wang Y, Zhang W, Wang X, Chen L, Wang S. BKM120 sensitizes BRCA-proficient triple negative breast cancer cells to olaparib through regulating FOXM1 and Exo1 expression. Sci Rep. 2021;11:4774.

53. Ibrahim YH, García-García C, Serra V, He L, Torres-Lockhart K, Prat A, et al. PI3K inhibition impairs BRCA1/2 expression and sensitizes BRCA-proficient triple-negative breast cancer to PARP inhibition. Cancer Discov. 2012;2:1036–1047.

54. Yap TA, Kristeleit R, Michalarea V, Pettitt SJ, Lim JSJ, Carreira S, et al. Phase I Trial of the PARP Inhibitor Olaparib and AKT Inhibitor Capivasertib in Patients with BRCA1/2- and Non-BRCA1/2-Mutant Cancers. Cancer Discov. 2020;10:1528–1543.

55. Abraham JE, Pinilla K, Dayimu A, Grybowicz L, Demiris N, Harvey C, et al. The PARTNER trial of neoadjuvant olaparib in triple-negative breast cancer. Nature. 2024;629:1142–1148.

56. Lee JM, Peer CJ, Yu M, Amable L, Gordon N, Annunziata CM, et al. Sequence-specific pharmacokinetic and pharmacodynamic phase I/Ib study of Olaparib tablets and carboplatin in women’s cancer. Clin Cancer Res. 2017;23:1397–1406.

57. Plana D, Palmer AC, Sorger PK. Independent drug action in combination therapy: implications for precision oncology. Cancer Discov. 2022;12:606–624.

