## Supplementary Figures and Tables for "PARP inhibition enhances exemestane efficacy in triple-negative breast cancer"

### Supplementary Information

**Supplementary Table S1: Details of the FDA-approved drugs set (166 compounds) obtained from the NCI.** NSC = Cancer Chemotherapy National Service Center number, USAN = United States Adopted Names and MW = molecular weight.

| PLATE | WELL ID | NSC | CAS | DRUG NAME (USAN) | MW (g/mol) |
|---|---|---|---|---|---|
| 4893 | A02 | 740 | 59-05-2 | Methotrexate | 454.44 |
| 4893 | B02 | 750 | 55-98-1 | Busulfan | 246.30 |
| 4893 | C02 | 752 | 154-42-7 | Thioguanine | 167.19 |

| | | | | | |
|---|---|---|---|---|---|
| 4893 | D02 | 755 | 50-44-2 | Mercaptopurine | 152.18 |
| 4893 | E02 | 762 | 55-86-7 | Mechlorethamine hydrochloride | 192.52 |
| 4893 | F02 | 1390 | 315-30-0 | Allopurinol | 136.11 |
| 4893 | G02 | 3053 | 50-76-0 | Dactinomycin | 1255.43 |
| 4893 | H02 | 3088 | 305-03-3 | Chlorambucil | 304.22 |
| 4893 | A03 | 6396 | 52-24-4 | Thiotepa | 189.22 |
| 4893 | B03 | 8806 | 3223-07-2 | Melphalan hydrochloride | 341.66 |
| 4893 | C03 | 9706 | 51-18-3 | Triethylenemelamine | 204.23 |
| 4893 | D03 | 13875 | 645-05-6 | Altretamine | 210.28 |
| 4893 | E03 | 18509 | 5451-09-2 | Aminolevulinic acid hydrochloride | 167.59 |
| 4893 | F03 | 19893 | 51-21-8 | Fluorouracil | 130.08 |
| 4893 | G03 | 24559 | 18378-89-7 | Plicamycin | 1085.16 |
| 4893 | H03 | 25154 | 54-91-1 | Pipobroman | 356.06 |
| 4893 | A04 | 26271 | 6055-19-2 | Cyclophosphamide | 261.09 |
| 4893 | B04 | 26980 | 50-07-7 | Mitomycin | 334.33 |
| 4893 | C04 | 27640 | 50-91-9 | Floxuridine | 246.19 |
| 4893 | D04 | 32065 | 127-07-1 | Hydroxyurea | 76.05 |
| 4893 | E04 | 34462 | 66-75-1 | Uracil mustard | 252.10 |
| 4893 | F04 | 38721 | 53-19-0 | Mitotane | 320.04 |
| 4893 | G04 | 45388 | 4342-03-4 | Dacarbazine | 182.18 |
| 4893 | H04 | 45923 | 298-81-7 | Methoxsalen | 216.19 |
| 4893 | A05 | 49842 | 143-67-9 | Vinblastine sulfate | 909.06 |
| 4893 | B05 | 63878 | 69-74-9 | Cytarabine hydrochloride | 279.70 |
| 4893 | C05 | 66847 | 50-35-1 | Thalidomide | 258.23 |
| 4893 | D05 | 67574 | 2068-78-2 | Vincristine sulfate | 923.04 |
| 4893 | E05 | 71423 | 595-33-5 | Megestrol acetate | 384.51 |
| 4893 | F05 | 75520 | 70-00-8 | Trifluridine | 296.20 |
| 4893 | G05 | 77213 | 366-70-1 | Procarbazine hydrochloride | 257.76 |
| 4893 | H05 | 79037 | 13010-47-4 | Lomustine | 233.70 |
| 4893 | A06 | 82151 | 23541-50-6 | Daunorubicin hydrochloride | 563.98 |
| 4893 | B06 | 85998 | 18883-66-4 | Streptozocin | 265.22 |
| 4893 | C06 | 92859 | 1327-53-3 | Arsenic trioxide | 197.84 |
| 4893 | D06 | 102816 | 320-67-2 | Azacitidine | 244.21 |
| 4893 | E06 | 105014 | 4291-63-8 | Cladribine | 285.69 |
| 4893 | F06 | 109724 | 3778-73-2 | Ifosfamide | 261.09 |
| 4893 | G06 | 119875 | 15663-27-1 | Cisplatin | 300.06 |
| 4893 | H06 | 122758 | 302-79-4 | Tretinoin | 300.44 |
| 4893 | A07 | 122819 | 29767-20-2 | Teniposide | 656.66 |
| 4893 | B07 | 123127 | 25316-40-9 | Doxorubicin hydrochloride | 579.99 |
| 4893 | C07 | 125066 | 9041-93-4 | Bleomycin sulfate | 1512.61 |
| 4893 | D07 | 125973 | 33069-62-4 | Paclitaxel | 853.92 |
| 4893 | E07 | 127716 | 2353-33-5 | Decitabine | 228.21 |
| 4893 | F07 | 138783 | 3543-75-7 | Bendamustine hydrochloride | 394.73 |
| 4893 | G07 | 141540 | 33419-42-0 | Etoposide | 588.56 |
| 4893 | H07 | 169780 | 24584-09-6 | Dexrazoxane | 268.27 |
| 4893 | A08 | 180973 | 54965-24-1 | Tamoxifen citrate | 563.65 |
| 4893 | B08 | 218321 | 53910-25-1 | Pentostatin | 268.27 |
| 4893 | C08 | 226080 | 53123-88-9 | Sirolimus | 914.18 |
| 4893 | D08 | 241240 | 41575-94-4 | Carboplatin | 371.25 |
| 4893 | E08 | 246131 | 56124-62-0 | Valrubicin | 723.64 |
| 4893 | F08 | 256439 | 57852-57-0 | Idarubicin hydrochloride | 533.96 |
| 4893 | G08 | 256942 | 56390-09-1 | Epirubicin hydrochloride | 579.99 |
| 4893 | H08 | 266046 | 61825-94-3 | Oxaliplatin | 397.29 |
| 4893 | A09 | 279836 | 65271-80-9 | Mitoxantrone | 444.49 |

| | | | | | |
|---|---|---|---|---|---|
| 4893 | B09 | 296961 | 20537-88-6 | Amifostine | 214.22 |
| 4893 | C09 | 312887 | 75607-67-9 | Fludarabine phosphate | 365.21 |
| 4893 | D09 | 362856 | 85622-93-1 | Temozolomide | 194.15 |
| 4893 | E09 | 369100 | 99011-02-6 | Imiquimod | 240.31 |
| 4893 | F09 | 409962 | 154-93-8 | Carmustine | 214.05 |
| 4893 | G09 | 606869 | 123318-82-1 | Clofarabine | 303.68 |
| 4893 | H09 | 608210 | 125317-39-7 | Vinorelbine tartrate | 1079.00 |
| 4893 | A10 | 609699 | 119413-54-6 | Topotecan hydrochloride | 457.91 |
| 4893 | B10 | 613327 | 122111-03-9 | Gemcitabine hydrochloride | 299.65 |
| 4893 | C10 | 616348 | 100286-90-6 | Irinotecan hydrochloride | 623.15 |
| 4893 | D10 | 628503 | 114977-28-5 | Docetaxel | 807.89 |
| 4893 | E10 | 683864 | 162635-04-3 | Temsirolimus | 1030.29 |
| 4893 | F10 | 701852 | 149647-78-9 | Vorinostat | 264.32 |
| 4893 | G10 | 702294 | 52205-73-9 | Estramustine phosphate sodium | 564.35 |
| 4893 | H10 | 712807 | 154361-50-9 | Capecitabine | 359.35 |
| 4893 | A11 | 713563 | 107868-30-4 | Exemestane | 296.40 |
| 4893 | B11 | 715055 | 184475-35-2 | Gefitinib | 446.90 |
| 4893 | C11 | 718781 | 183319-69-9 | Erlotinib hydrochloride | 429.90 |
| 4893 | D11 | 719276 | 129453-61-8 | Fulvestrant | 606.75 |
| 4893 | E11 | 719344 | 120511-73-1 | Anastrozole | 293.37 |
| 4893 | F11 | 719345 | 112809-51-5 | Letrozole | 285.30 |
| 4893 | G11 | 719627 | 169590-42-5 | Celecoxib | 381.37 |
| 4893 | H11 | 721517 | 118072-93-8 | Zoledronic acid | 272.09 |
| 4894 | A02 | 732517 | 863127-77-9 | Dasatinib | 488.01 |
| 4894 | B02 | 733504 | 159351-69-6 | Everolimus | 958.24 |
| 4894 | C02 | 737754 | 635702-64-6 | Pazopanib hydrochloride | 473.98 |
| 4894 | D02 | 741078 | 606143-52-6 | Selumetinib | 457.68 |
| 4894 | E02 | 743414 | 152459-95-5 | Imatinib | 493.61 |
| 4894 | F02 | 745750 | 231277-92-2 | Lapatinib | 581.06 |
| 4894 | G02 | 747599 | 641571-10-0 | Nilotinib | 529.51 |
| 4894 | H02 | 747971 | 284461-73-0 | Sorafenib | 464.82 |
| 4894 | A03 | 747972 | 191732-72-6 | Lenalidomide | 259.26 |
| 4894 | B03 | 747973 | 219989-84-1 | Ixabepilone | 506.70 |
| 4894 | C03 | 747974 | 84449-90-1 | Raloxifene | 473.59 |
| 4894 | D03 | 749226 | 154229-19-3 | Abiraterone | 349.51 |
| 4894 | E03 | 750690 | 557795-19-4 | Sunitinib | 398.47 |
| 4894 | F03 | 750691 | 439081-18-2 | Afatinib | 485.94 |
| 4894 | G03 | 753686 | 763113-22-0 | Olaparib | 434.46 |
| 4894 | H03 | 754143 | 128517-07-7 | Romidepsin | 540.69 |
| 4894 | A04 | 754230 | 146464-95-1 | Pralatrexate | 477.48 |
| 4894 | B04 | 754355 | 1038915-60-4 | Niraparib hydrochloride | 356.85 |
| 4894 | C04 | 755384 | 357166-30-4 | Pemetrexed, Disodium salt, Heptahydrate | 471.38 |
| 4894 | D04 | 755605 | 915087-33-1 | Enzalutamide | 464.42 |
| 4894 | E04 | 755980 | 417716-92-8 | Lenvatinib | 426.86 |
| 4894 | F04 | 755985 | 121032-29-9 | Nelarabine | 297.27 |
| 4894 | G04 | 755986 | 879085-55-9 | Vismodegib | 421.30 |
| 4894 | H04 | 756644 | 459868-92-9 | Rucaparib phosphate | 421.36 |
| 4894 | A05 | 756645 | 877399-52-5 | Crizotinib | 450.34 |
| 4894 | B05 | 756655 | 179324-69-7 | Bortezomib | 384.24 |
| 4894 | C05 | 757439 | 698387-09-6 | Neratinib | 557.05 |
| 4894 | D05 | 757441 | 319460-85-0 | Axitinib | 386.47 |
| 4894 | E05 | 758246 | 871700-17-3 | Trametinib | 615.40 |
| 4894 | F05 | 758247 | 571190-30-2 | Palbociclib | 447.54 |
| 4894 | G05 | 758252 | 868540-17-4 | Carfilzomib | 719.92 |

| | | | | | |
|---|---|---|---|---|---|
| 4894 | H05 | 758253 | 26833-87-4 | Omacetaxine mepesuccinate | 545.63 |
| 4894 | A06 | 758254 | 1201902-80-8 | Ixazomib citrate | 517.13 |
| 4894 | B06 | 758487 | 943319-70-8 | Ponatinib | 532.55 |
| 4894 | C06 | 758774 | 414864-00-9 | Belinostat | 318.35 |
| 4894 | D06 | 759224 | 870281-82-6 | Idelalisib | 415.42 |
| 4894 | E06 | 760766 | 443913-73-3 | Vandetanib | 475.36 |
| 4894 | F06 | 761068 | 849217-68-1 | Cabozantinib | 501.51 |
| 4894 | G06 | 761190 | 404950-80-7 | Panobinostat | 349.43 |
| 4894 | H06 | 761385 | 956697-53-3 | Erismodegib | 485.49 |
| 4894 | A07 | 761388 | 110078-46-1 | Plerixafor | 502.79 |
| 4894 | B07 | 761431 | 1029872-54-5 | Vemurafenib | 489.91 |
| 4894 | C07 | 761432 | 183133-96-2 | Cabazitaxel | 835.94 |
| 4894 | D07 | 761910 | 936563-96-1 | Ibrutinib | 440.50 |
| 4894 | E07 | 763371 | 941678-49-5 | Ruxolitinib | 306.36 |
| 4894 | F07 | 763932 | 755037-03-7 | Regorafenib | 482.80 |
| 4894 | G07 | 764040 | 1256580-46-7 | Alectinib | 482.62 |
| 4894 | H07 | 764042 | 606143-89-9 | Binimetinib | 441.22 |
| 4894 | A08 | 764134 | 1195768-06-9 | Dabrafenib mesylate | 615.65 |
| 4894 | B08 | 764581 | 937263-43-9 | ARRY-380 | 480.52 |
| 4894 | C08 | 765694 | 380843-75-4 | Bosutinib | 530.45 |
| 4894 | D08 | 765888 | 1110813-31-4 | Dacomitinib | 469.94 |
| 4894 | E08 | 765974 | 1217486-61-7 | Alpelisib | 441.47 |
| 4894 | F08 | 766270 | 1257044-40-8 | Venetoclax | 868.45 |
| 4894 | G08 | 767125 | 1207456-01-6 | Talazoparib | 380.35 |
| 4894 | H08 | 767600 | 936091-26-8 | Fedratinib | 524.67 |
| 4894 | A09 | 768068 | 934660-93-2 | Cobimetinib | 531.32 |
| 4894 | B09 | 768073 | 1231929-97-7 | Abemaciclib | 506.59 |
| 4894 | C09 | 771649 | 956104-40-8 | Apalutamide | 477.42 |
| 4894 | D09 | 772469 | 1201438-56-3 | Duvelisib | 416.87 |
| 4894 | E09 | 774769 | 1108743-60-7 | Entrectinib | 560.63 |
| 4894 | F09 | 775351 | 19171-19-8 | Pomalidomide | 273.25 |
| 4894 | G09 | 775772 | 1095173-27-5 | Glasdegib | 374.43 |
| 4894 | H09 | 776422 | 1032900-25-6 | Ceritinib | 558.14 |
| 4894 | A10 | 777109 | 1403254-99-8 | Tazemetostat | 572.73 |
| 4894 | B10 | 777878 | 1029712-80-8 | Capmatinib | 412.41 |
| 4894 | C10 | 778304 | 1269440-17-6 | Encorafenib | 540.01 |
| 4894 | D10 | 778909 | 1211441-98-3 | Ribociclib | 434.54 |
| 4894 | E10 | 779217 | 1421373-65-0 | Osimertinib | 499.61 |
| 4894 | F10 | 780108 | 1454846-35-5 | Lorlatinib | 406.41 |
| 4894 | G10 | 780203 | 1393477-72-9 | Selinexor | 443.30 |
| 4894 | H10 | 781556 | 1346242-81-6 | Erdafitinib | 446.54 |
| 4894 | A11 | 785570 | 1223403-58-4 | Larotrectinib | 428.43 |
| 4894 | B11 | 787457 | 1197953-54-0 | Brigatinib | 584.10 |
| 4894 | C11 | 787846 | 1254053-43-4 | Gilteritinib | 552.71 |
| 4894 | D11 | 788120 | 1446502-11-9 | Enasidenib | 473.35 |
| 4894 | E11 | 788948 | 4105-38-8 | Uridine triacetate | 370.32 |
| 4894 | F11 | 789102 | 1448347-49-6 | Ivosidenib | 582.96 |
| 4894 | G11 | 789300 | 1029044-16-3 | Pexidartinib | 417.81 |
| 4894 | H11 | 791164 | 1420477-60-6 | Acalabrutinib | 465.41 |
| 4895 | A02 | 801082 | 1703793-34-3 | Avapritinib | 498.55 |
| 4895 | B02 | 816437 | N/A | Copanlisib tris-HCl | 589.90 |
| 4895 | C02 | 816556 | 1513857-77-6 | Pemigatinib | 487.49 |
| 4895 | D02 | 818434 | 2152628-33-4 | Selpercatinib | 525.60 |
| 4895 | E02 | 823807 | 1691249-45-2 | Zanubrutinib | 471.55 |

| | | | | | |
|---|---|---|---|---|---|
| 4895 | F02 | 825331 | 1297538-32-9 | Darolutamide | 398.84 |

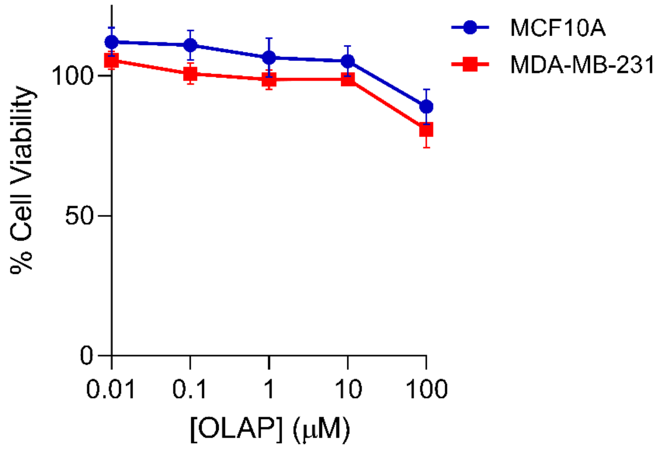


**Supplementary Fig. S1: Cell viability of MCF10A normal breast cells and MDA-MB-231 TNBC cancer cell lines following treatment with concentration gradient of Olaparib for 24 h, as determined by MTT assay.** Data were expressed as mean ± SD of three independent experiments.

**Supplementary Table S2: Combination indices (CIs) for the combinations of the library compound with Olaparib in MDA-MB-231 cells for 24 h treatment.** Sub-cytotoxic concentration of Olaparib (10 µM) was used in the combination treatment.

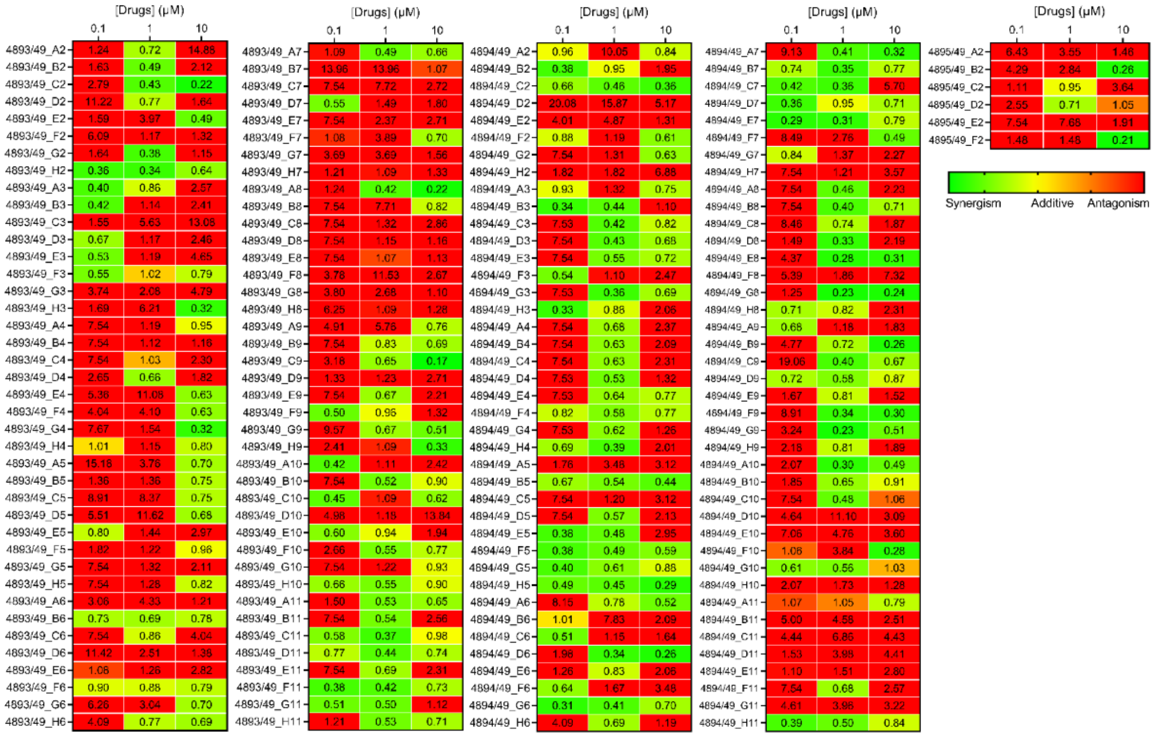


**Supplementary Table S3: $IC_{50}$ values of each compound alone or in combination with Olaparib (10 µM) in MDA-MB-231 cells, 24 h treatment, as determined from MTT data**. Data expressed as mean ± SD of two independent experiments. Fold shift = $IC_{50}$ (- OLAP)/$IC_{50}$ (+ OLAP). ND = not determined.

| Compound | | 24 h $IC_{50}$ (µM) | | Fold Change |
|---|---|---|---|---|
| | | - 10 µM OLAP | + 10 µM OLAP | |
| 4893/49_H2 | Chlorambucil | >100 | 60.0 ± 12.4 | > 1.6 |
| 4893/49_A8 | Tamoxifen Citrate | >100 | 15.4 ± 0.9 | > 6.4 |
| 4893/49_C9 | Fludarabine Phosphate | >100 | 0.5 ± 0.1 | > 200 |

| | | | | |
|---|---|---|---|---|
| 4893/49_A11 | Exemestane | >100 | 16.8 ± 12.5 | > 5.9 |
| 4893/49_H11 | Zoledronic acid | >100 | 51.9 ± 0.9 | > 1.9 |
| 4894/49_D3 | Abiraterone | >100 | 21.8 ± 10.4 | > 4.5 |
| 4894/49_H5 | Omacetaxine mepesuccinate | 1.6 ± 1.0 | 0.2 ± 0.1 | 8 |
| 4894/49_G6 | Panobinostat | 4.1 ± 2.8 | 0.1 ± 0.0 | 41 |
| 4894/49_A7 | Plerixafor | >100 | 66.6 ± 11.6 | > 1.5 |
| 4894/49_H11 | Acalabrutinib | >100 | >100 | ND |

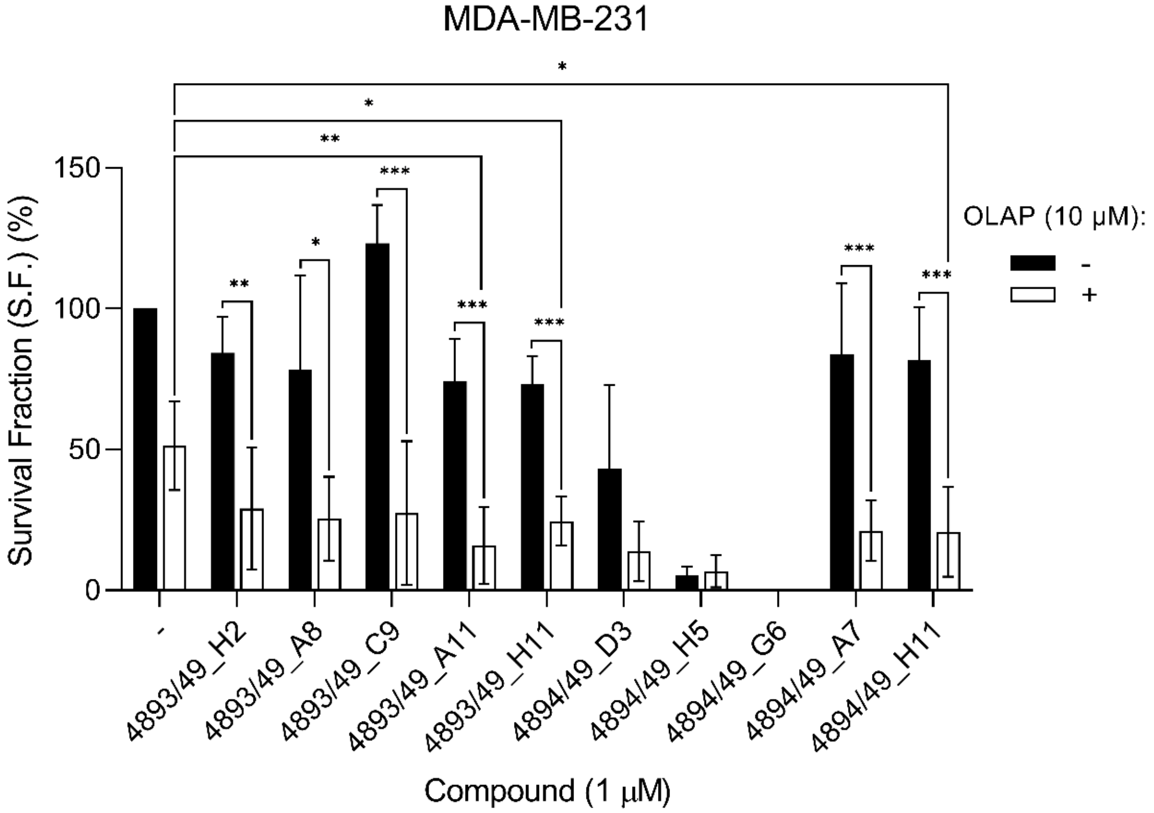


**Supplementary Fig. S2: Clonogenic survival assay of MDA-MB-231 cells treated with the stated single-agent (1 µM), Olaparib (10 µM) or both for 24 h.** Data were expressed as mean ± SD of four independent experiments (n = 4). *$P$ < 0.05, **$P$ < 0.01 and ***$P$ < 0.001 compared to single agent-treated groups by ANOVA.

**Supplementary Table S4 Detail of the ten hit compounds including their drug classes, human safety profiles of these drugs and their PARPi combinations status.** NA = Not Available.

| Compound | Drug class | Human safety profiles | PARPi combination |
|---|---|---|---|
| Chlorambucil | DNA alkylating agent | Severe toxicity | NA |
| Tamoxifen citrate | a selective oestrogen receptor modulator | NA | [1] |
| Fludarabine phosphate | a purine analogue antimetabolite | Severe toxicity | NA |
| Exemestane | aromatase inhibitor | NA | NA |
| Zoledronic acid | bisphosphonate | NA | NA |
| Abiraterone | antiandrogen | NA | [2] |
| Omacetaxine mepesuccinate | cephalotaxine | NA | NA |
| Panobinostat | non-selective histone deacetylase inhibitor | Severe toxicity | NA |
| Plerixafor | a selective chemokine receptor (CXCR4) antagonist | NA | NA |
| Acalabrutinib | a Bruton tyrosine kinase inhibitor | NA | NA |

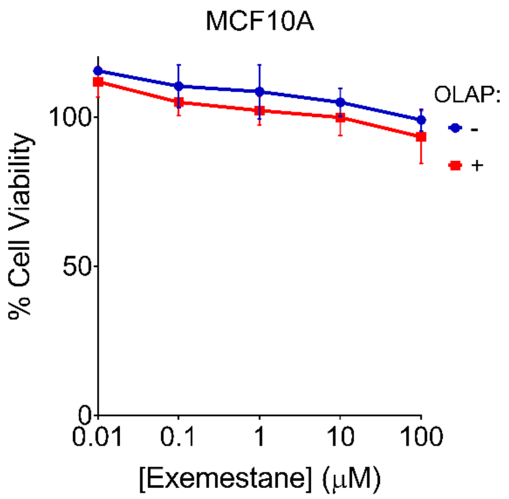


**Supplementary Fig. S3: Cell viability of MCF10A normal breast cells following treatment with concentration gradients of Exemestane, with and without 10 μM Olaparib (72 h treatment), as determined by MTT assay.**

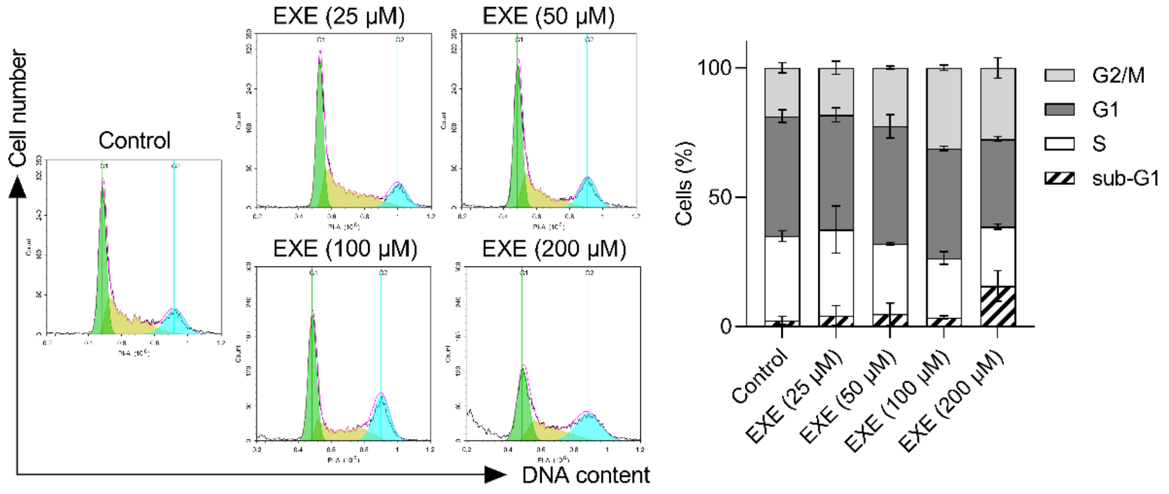


**Supplementary Fig. S4: Cell cycle distribution following 24 h treatment with Exemestane (25, 50, 100 and 200 µM, as determined by PI staining and flow cytometry.** Left, representative histograms, right, quantification of cell cycle phase.

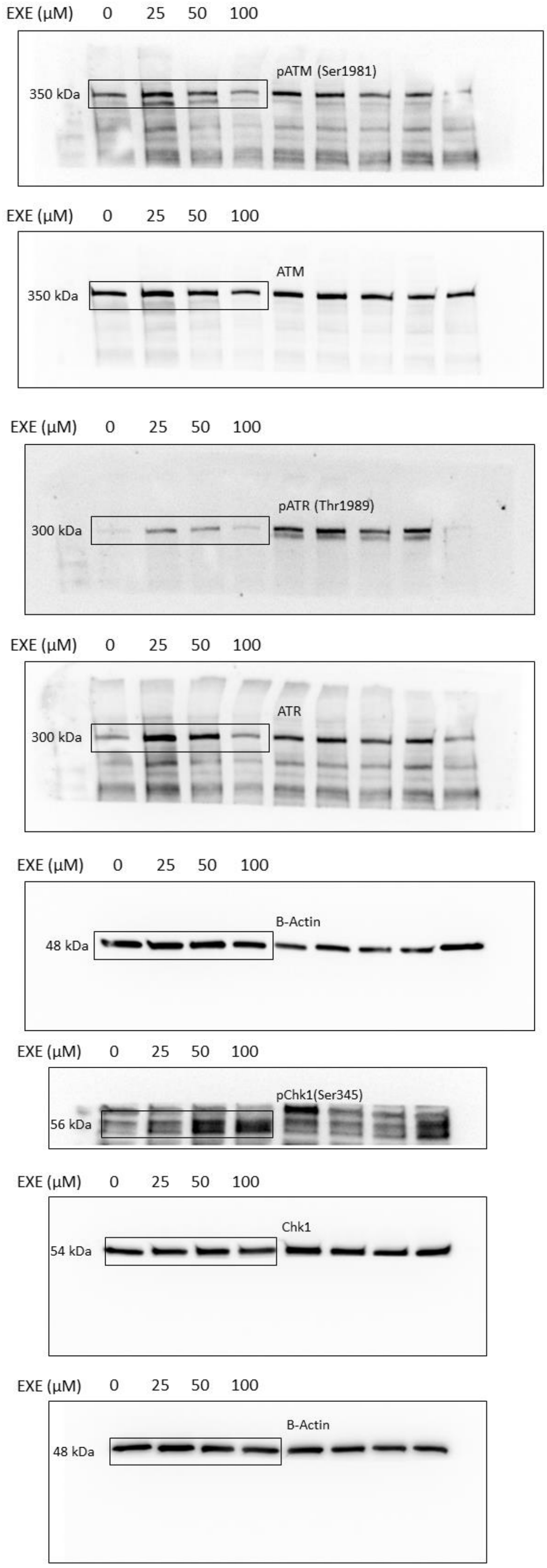


**Supplementary Fig. S5: Uncropped blot images from Figure 3 with molecular weight (kDa) of the protein of interest on the left.**

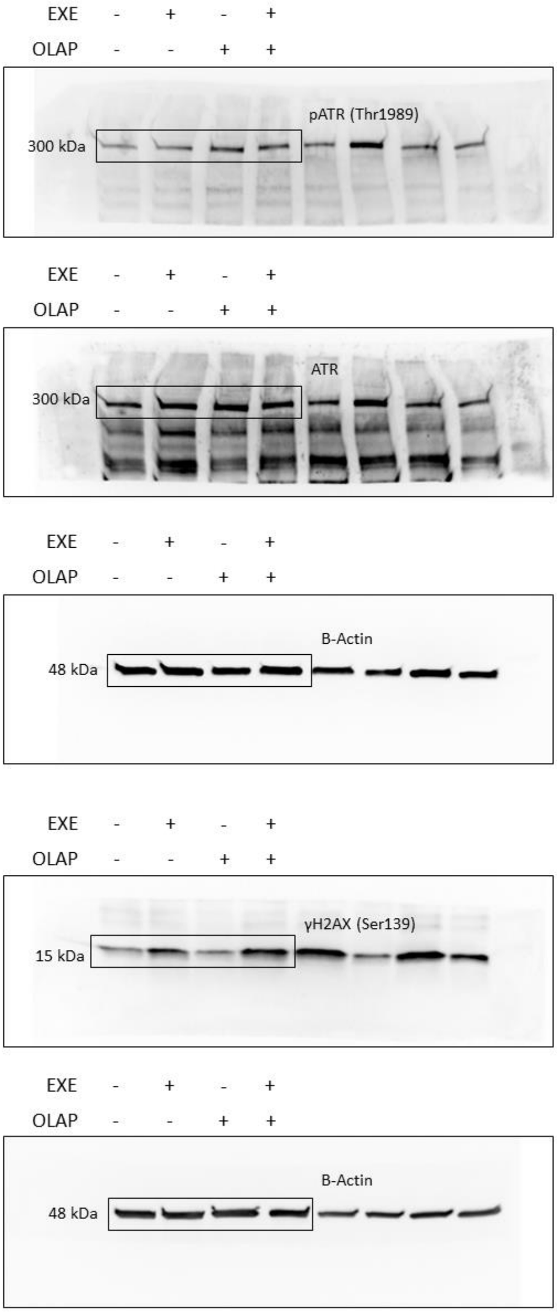


**Supplementary Fig. S6: Uncropped blot images from Figure 4 with molecular weight (kDa) of the protein of interest on the left.**

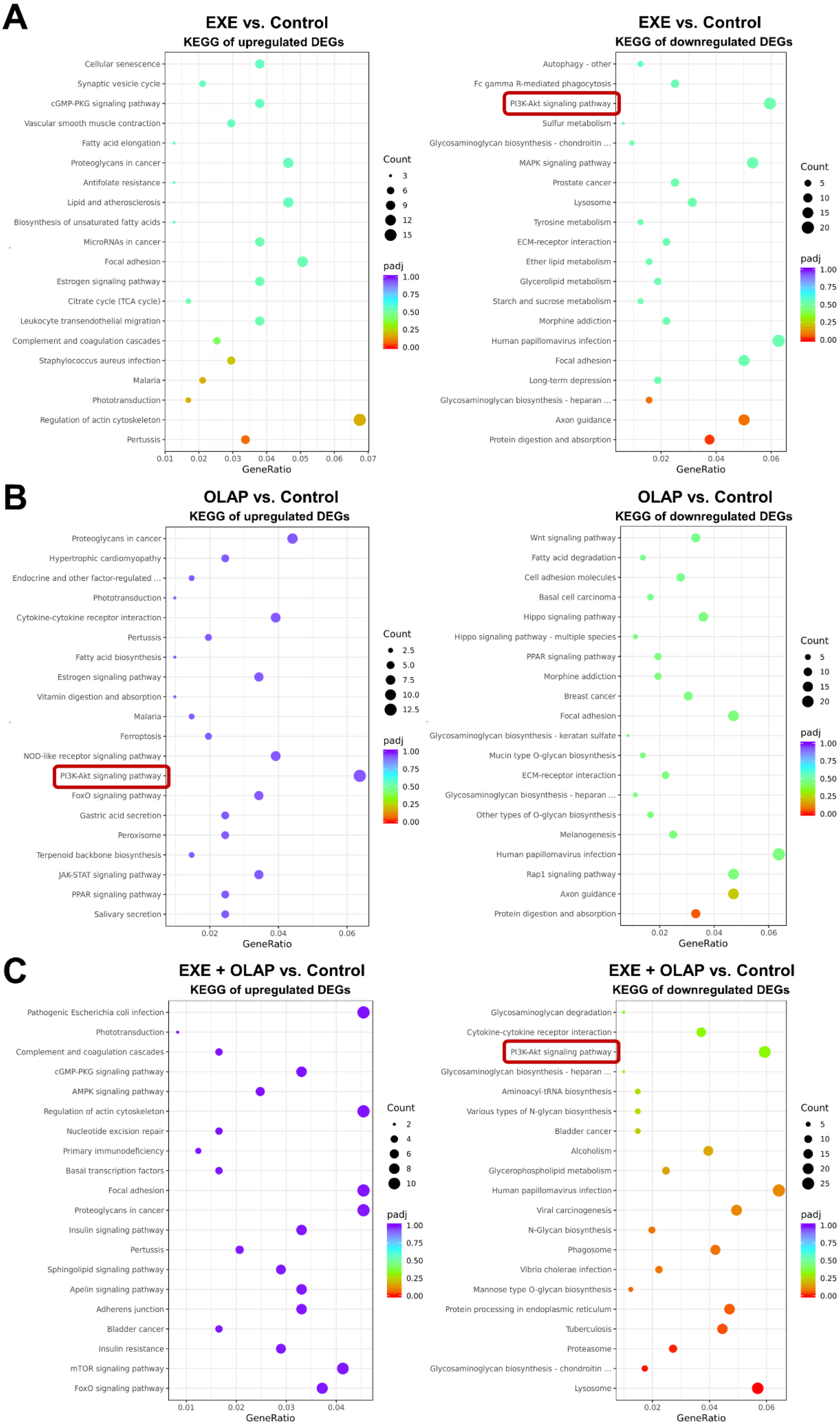
A
EXE vs. Control
KEGG of upregulated DEGs
Cellular senescence
Synaptic vesicle cycle
cGMP-PKG signaling pathway
Vascular smooth muscle contraction
Fatty acid elongation
Proteoglycans in cancer
Antifolate resistance
Lipid and atherosclerosis
Biosynthesis of unsaturated fatty acids
MicroRNAs in cancer
Focal adhesion
Estrogen signaling pathway
Citrate cycle (TCA cycle)
Leukocyte transendothelial migration
Complement and coagulation cascades
Staphylococcus aureus infection
Malaria
Phototransduction
Regulation of actin cytoskeleton
Pertussis
0.01 0.02 0.03 0.04 0.05 0.06 0.07
GeneRatio
Count
3
6
9
12
15
padj
1.00
0.75
0.50
0.25
0.00
EXE vs. Control
KEGG of downregulated DEGs
Autophagy - other
Fc gamma R-mediated phagocytosis
PI3K-Akt signaling pathway
Sulfur metabolism
Glycosaminoglycan biosynthesis - chondroitin ...
MAPK signaling pathway
Prostate cancer
Lysosome
Tyrosine metabolism
ECM-receptor interaction
Ether lipid metabolism
Glycerolipid metabolism
Starch and sucrose metabolism
Morphine addiction
Human papillomavirus infection
Focal adhesion
Long-term depression
Glycosaminoglycan biosynthesis - heparan ...
Axon guidance
Protein digestion and absorption
0.02 0.04 0.06
GeneRatio
Count
5
10
15
20
padj
1.00
0.75
0.50
0.25
0.00
B
OLAP vs. Control
KEGG of upregulated DEGs
Proteoglycans in cancer
Hypertrophic cardiomyopathy
Endocrine and other factor-regulated ...
Phototransduction
Cytokine-cytokine receptor interaction
Pertussis
Fatty acid biosynthesis
Estrogen signaling pathway
Vitamin digestion and absorption
Malaria
Ferroptosis
NOD-like receptor signaling pathway
PI3K-Akt signaling pathway
FoxO signaling pathway
Gastric acid secretion
Peroxisome
Terpenoid backbone biosynthesis
JAK-STAT signaling pathway
PPAR signaling pathway
Salivary secretion
0.02 0.04 0.06
GeneRatio
Count
2.5
5.0
7.5
10.0
12.5
padj
1.00
0.75
0.50
0.25
0.00
OLAP vs. Control
KEGG of downregulated DEGs
Wnt signaling pathway
Fatty acid degradation
Cell adhesion molecules
Basal cell carcinoma
Hippo signaling pathway
Hippo signaling pathway - multiple species
PPAR signaling pathway
Morphine addiction
Breast cancer
Focal adhesion
Glycosaminoglycan biosynthesis - keratan sulfate
Mucin type O-glycan biosynthesis
ECM-receptor interaction
Glycosaminoglycan biosynthesis - heparan ...
Other types of O-glycan biosynthesis
Melanogenesis
Human papillomavirus infection
Rap1 signaling pathway
Axon guidance
Protein digestion and absorption
0.02 0.04 0.06
GeneRatio
Count
5
10
15
20
padj
1.00
0.75
0.50
0.25
0.00
C
EXE + OLAP vs. Control
KEGG of upregulated DEGs
Pathogenic Escherichia coli infection
Phototransduction
Complement and coagulation cascades
cGMP-PKG signaling pathway
AMPK signaling pathway
Regulation of actin cytoskeleton
Nucleotide excision repair
Primary immunodeficiency
Basal transcription factors
Focal adhesion
Proteoglycans in cancer
Insulin signaling pathway
Pertussis
Sphingolipid signaling pathway
Apelin signaling pathway
Adherens junction
Bladder cancer
Insulin resistance
mTOR signaling pathway
FoxO signaling pathway
0.01 0.02 0.03 0.04
GeneRatio
Count
2
4
6
8
10
padj
1.00
0.75
0.50
0.25
0.00
EXE + OLAP vs. Control
KEGG of downregulated DEGs
Glycosaminoglycan degradation
Cytokine-cytokine receptor interaction
PI3K-Akt signaling pathway
Glycosaminoglycan biosynthesis - heparan ...
Aminoacyl-tRNA biosynthesis
Various types of N-glycan biosynthesis
Bladder cancer
Alcoholism
Glycerophospholipid metabolism
Human papillomavirus infection
Viral carcinogenesis
N-Glycan biosynthesis
Phagosome
Vibrio cholerae infection
Mannose type O-glycan biosynthesis
Protein processing in endoplasmic reticulum
Tuberculosis
Proteasome
Glycosaminoglycan biosynthesis - chondroitin ...
Lysosome
0.02 0.04 0.06
GeneRatio
Count
5
10
15
20
25
padj
1.00
0.75
0.50
0.25
0.00

**Supplementary Fig. S7: KEGG pathway enrichment analysis of DEGs in transcriptomes of (A) EXE-, (B) OLAP-, and (C) EXE+OLAP-treated groups in comparison to control group.** The top 20 significantly enriched pathways are presented in order of enrichment score. Dot size represents gene count, and colour represents the *P* value. $P < 0.05$ is considered statistically significant.

**REFERENCES**

1. Plummer R, Verheul HM, De Vos F, Leunen K, Molife LR, Rolfo C , et al. Pharmacokinetic Effects and Safety of Olaparib Administered with Endocrine Therapy: A Phase I Study in Patients with Advanced Solid Tumours. Adv Ther. 2018;35:1945-1964.

2. Clarke NW, Armstrong AJ, Thiery-Vuillemin A, Oya M, Shore N, Loredo E , et al. Abiraterone and Olaparib for Metastatic Castration-Resistant Prostate Cancer. NEJM Evidence. 2022;1:EVIDoa2200043.
